# Endothelial NRP2 influences angiogenesis by regulating actin pattern development and α5-integrin-*p*-FAK complex recruitment to assembling adhesion sites

**DOI:** 10.1101/2021.01.15.426835

**Authors:** Christopher J Benwell, James AGE Taylor, Stephen D Robinson

## Abstract

The ability to form a variety of cell-matrix connections is crucial for angiogenesis to take place. Without stable anchorage to the extracellular matrix (ECM), endothelial cells (ECs) are unable to sense, integrate and disseminate growth factor stimulated responses that drive growth of a vascular bed. Neuropilin-2 (NRP2) is a widely expressed membrane-bound multifunctional non-tyrosine kinase receptor, that has previously been implicated in influencing cell adhesion and migration by interacting with α5-integrin and regulating adhesion turnover. α5-integrin, and its ECM ligand fibronectin (FN) are both known to be upregulated during the formation of neo-vasculature. Despite being descriptively annotated as a candidate biomarker for aggressive cancer phenotypes, the EC-specific roles for NRP2 during developmental and pathological angiogenesis remain unexplored. The data reported here support a model whereby NRP2 actively promotes EC adhesion and migration by regulating dynamic cytoskeletal remodelling and by stimulating Rab11-dependent recycling of α5-integrin-*p*-FAK complexes to newly assembling adhesion sites. Furthermore, temporal depletion of EC-NRP2 *in vivo* impairs primary tumour growth by disrupting vessel formation. We also demonstrate that EC-NRP2 is required for normal postnatal retinal vascular development, specifically by regulating cell-matrix adhesion. Upon loss of endothelial NRP2, vascular outgrowth from the optic nerve during superficial plexus formation is disrupted, likely due to reduced FAK phosphorylation within sprouting tip cells.

## Introduction

Angiogenesis, the formation of neo-vasculature from pre-existing blood vessels is essential to moderate the hypoxic stress that arises in tumour microenvironments (Hanahan and Weinberg, 2011). Sustained primary tumour growth therefore relies on the protracted stimulation of neighbouring endothelial cells (ECs) to activate appropriate growth factor receptors such as vascular endothelial growth factor (VEGF) receptor-2 (VEGFR2) on their cell surface. The ensuing proliferation and integrin-dependent migration of ECs towards the angiogenic stimulus then results in the formation of a dense network of highly permeable blood vessels to re-perfuse the hypoxic tumour niche (Robinson and Hodivala-Dilke, 2011)

EC migration relies on the cell’s ability to propagate integrin outside-in signalling to the extracellular space (Woodside, Liu and Ginsberg, 2001; Calderwood, 2004; Pan *et al.*, 2016). Upon ligand binding, integrins mediate the recruitment of core cytosolic scaffolding proteins at the sites of adhesion to the extracellular matrix (ECM) to form focal adhesions (FAs) (Hynes, 2002; Hu and Luo, 2013; De Pascalis and Etienne-Manneville, 2017; Buskermolen, Kurniawan and Bouten, 2018). Additional recruitment and clustering of protein kinases such as focal adhesion kinase (FAK) at FAs following integrin-mediated adhesion to the ECM subsequently enables integrin receptors to overcome the limits imposed by the absence of enzymatic or kinase activity in their cytoplasmic tails, allowing for the potentiated activation of numerous signalling cascades (Srichai and Zent, 2010). An essential downstream effector for directional cell migration, FAK also indirectly mediates the activation and translocation of Rac1 to FAs, which promotes the nucleation and polymerisation of actin stress fibres. This enables for the extension and protrusion of lamellipodia and filopodia at the leading edge to drive cell spreading and migration (Chang *et al.*, 2007; Hu and Luo, 2013; Sadok and Marshall, 2014). Dynamic remodelling of the actin cytoskeleton is also coupled to the rapid disassembly and re-assembly of nascent FAs, which utilise actin highways to coordinate integrin transport and trafficking (Goley and Welch, 2006; Kaksonen, Toret and Drubin, 2006; Valdembri *et al.*, 2009; Sundararaman *et al.*, 2020).

Neuropilin-2 (NRP2) is a single pass transmembrane receptor whose upregulation is consistent with an accelerated cancer progression in a number of cell types [e.g., neuroblastomas (Fakhari *et al.*, 2002), non-small cell lung carcinoma [NSCLC] (Kawakami *et al.*, 2002), human prostate carcinoma, melanoma (Bielenberg *et al.*, 2004), lung cancer (Lantuéjoul *et al.*, 2003), myeloid leukaemia (Vales *et al.*, 2007), breast cancer (Bachelder *et al.*, 2001) and pancreatic cancer (Fukahi *et al.*, 2004)]. As a selective co-receptor for members of the VEGF family of growth factors, the function of NRP2 has also been implicated in influencing EC adhesion, migration and permeability during angiogenesis, under both physiological and pathological conditions (Favier *et al.*, 2006; Plein, Fantin and Ruhrberg, 2014; Alghamdi *et al.*, 2020). NRP2 has since emerged as a promising candidate for targeted therapy against tumorigenesis, however the mechanisms by which it integrates and disseminates ECM and growth factor signals to coordinate EC responses during angiogenesis is as yet unclear.

We have previously shown that endothelial NRP2 drives a VEGF-independent, Rac1-mediated mechanism of cell adhesion and migration over fibronectin (FN) matrices and regulates FA turnover by promoting basal α5-integrin recycling (Alghamdi *et al.*, 2020). Here we show that NRP2 regulates VASP-mediated control of actin pattern development to support the protrusion of dynamic lamellipodia at the leading edge of migrating cells. We also elucidate the existence of a NRP2-regulated phospho-FAK (*p*-FAK)-Rab11 trafficking axis, siRNA-mediated depletion of NRP2 in vascular ECs dampening the activation and recruitment of *p*-FAK to α5-integrin containing adhesions. We build on our previous work by demonstrating that the reduction in FA turnover rate elicited by NRP2 silencing leads to an accelerated growth of tensin-1 positive fibrillar adhesions, which in turn promotes FN fibrillogenesis. Finally, by utilising a mouse model that expresses a tamoxifen-inducible Cre-recombinase under the control of a PDGFb promoter, we examine the effects of specifically and temporally depleting endothelial NRP2 *in vivo.* To this end we demonstrate that loss of endothelial NRP2 inhibits tumorigenesis, and transiently impairs physiological vascularisation of developing postnatal retinae. This work highlights the advantages of employing endothelial-specific targeting therapies against pathological angiogenesis, in addition to providing further evidence that NRP2 is a promising anti-angiogenic target to impair primary tumour growth.

## Results

### VASP-mediated actin pattern development is sensitive to the loss of NRP2

Angiogenesis relies on the ability of ECs to sense, integrate and disseminate signals they receive from the ECM and from secreted growth factors in order to adhere and migrate towards the angiogenic stimulus (Clark *et al.*, 1982; Ingber, 1990). We have previously used multiple specific NRP2 siRNAs in both immortalised and primary mouse-lung microvascular ECs (mLMECs) to identify endothelial NRP2 as a key regulator of EC adhesion and migration by promoting FA turnover and Rac1 activation, independent of VEGF-stimulation. Label-free quantitative mass spectrometry was additionally utilised to elucidate candidate NRP2 interactions with several endocytic and recycling-associated trafficking proteins (Alghamdi *et al.*, 2020).

Migrating cells rely on the stimulation of numerous canonical signalling networks that converge on the organised remodelling of the cytoskeleton (Bökel and Brown, 2002). Actin is the major cytoskeletal element in ECs, and continuously undergoes stress fibre remodelling into dynamic filopodial and lamellipodial protrusions to maintain the cell in a motile state (Hu and Luo, 2013; Sadok and Marshall, 2014; Nader, Ezratty and Gundersen, 2016). Given NRP2’s regulation of Rac1 activity, we decided to consider its involvement during actin cytoskeletal remodelling in mLMECs. Luo et al., recently reported that pancreatic neuroendocrine tumour (PNET)-associated human umbilical vein ECs (HUVECs) ectopically overexpressing NRP2 formed larger actin-rich lamellipodial protrusions at the cell periphery compared to control ECs. This was subsequently shown to result from an upregulation of cofilin activity, propagating increased rates of actin polymerisation at the leading edges of the cell (Luo *et al.*, 2020). To investigate whether NRP2 is dispensable for actin cytoskeletal remodelling during initial adhesion to the ECM, we allowed control (*si*Ctrl) and NRP2-siRNA (*si*NRP2)-treated ECs to adhere to FN for 90 minutes before fixation and stained for filamentous actin (Fig.1A). Whilst Ctrl ECs formed long linear stress fibres ending in actin-rich lamellipodial protrusions, NRP2 siRNA-treated ECs appeared more circular, with visibly fewer linear stress fibres, protrusions and actin microspikes (Fig.1B). A similar observation was reported by Fantin et al., following NRP1 knockdown in hMVECs, delineating a role for NRP1 in the regulation of Rac1-mediated actin pattern development (Raimondi *et al.*, 2014; Fantin *et al.*, 2015).

**Figure 1:**
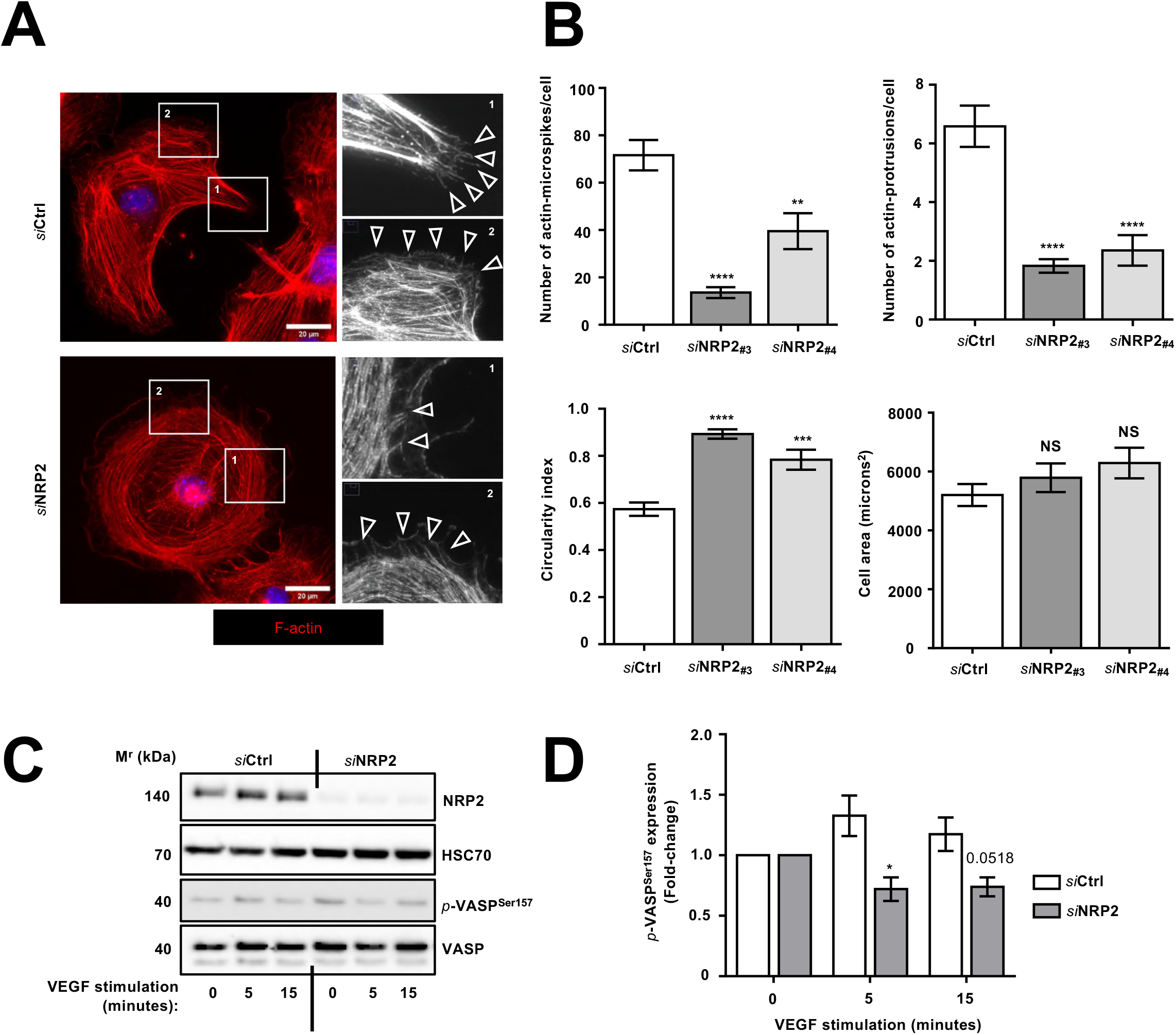
VASP-mediated actin pattern development and FAK phosphorylation are sensitive to the loss of NRP2. **A)** siRNA-transfected ECs were seeded onto FN coated coverslips and allowed to adhere for 90 minutes at 37°C and 5% CO_2_ before being fixed. ECs were then stained for filamentous actin using phalloidin Alexa Flour-568. Images were taken using a Zeiss AxioImager M2 microscope at 63X magnification. Arrowheads point to actin microspikes at cell periphery. **B)** Quantification of number of actin microspikes and actin protrusions/cell, cell circularity calculated as 4PiArea/Perimeter^2^, and cell area (microns^2^) in Ctrl siRNA and NRP2 siRNA-treated ECs. Two separate NRP2-targeting siRNAs (#3 and #4) were tested to confirm results. Error bars show ±SEM; n≥15 cells/siRNA-treated group. Asterisks indicate statistical significance from an unpaired two-tailed *t*-test. **C**) siRNA-transfected ECs were seeded onto FN and allowed to adhere for 48 hours at 37°C and 5% CO_2_. ECs were subsequently starved in serum free medium and stimulated with VEGF (30 ng/ml) for the indicated time. EC extracts were immunoblotted using antibodies against NRP2, HSC70, phospho-VASP (*p*-VASP) (Ser157) and VASP. Panel shows representative western blot image. **D**) Densitometric quantification of mean *p*-VASP (Ser157) band intensities normalised against total VASP expression and obtained using ImageJ™. Error bars show ± SEM; N=3 independent experiments. Asterisks indicate statistical significance from an unpaired two-tailed *t*-test (compared to *si*Ctrl).

To further elucidate the mechanisms by which NRP2 controls the remodelling of the actin cytoskeleton, we re-examined our label-free mass spectrometry data, originally performed to take an unbiased approach to identifying candidate proteins associating with NRP2 in mLMECs (Alghamdi *et al.*, 2020). This re-analysis revealed a large number of proteins known to be involved in regulating the actin cytoskeleton. Of note, this included those known to stabilise the spectrin actin network such as tropomyosin-1, members of the Arp2/3 complex, including cortactin, which regulates branch nucleation and polymerisation, and proteins involved in coordinating stress fibre development, such as VASP, cofilin-1, fascin, and TRIO-binding protein (Woo and Fowler, 1994; Yamakita *et al.*, 1996; Uruno *et al.*, 2001; Luo *et al.*, 2020). Unlike cofilin proteins, VASP functions as a potent regulator of both radial fibre formation and chiral actin pattern development, its depletion in fibroblasts preventing the transition from initial actin-rich peripheral rings to long linear stress fibres that support growth of dynamic protrusions (Benz *et al.*, 2009; Jalal *et al.*, 2019). As NRP2 depletion appeared to impair the formation of linear stress fibres, we next investigated whether NRP2 regulates VASP activity by monitoring total and phosphorylated VASP expression in the presence or absence of VEGF stimulation. Whilst total VASP expression was unaltered following NRP2 siRNA treatment, NRP2 depletion elicited a reduction in VASP phosphorylation at residue Ser157 at both 5 and 15 minutes post VEGF stimulation (Fig.1C-D). Phosphorylation at Ser157 provides a signal for membrane or leading-edge localisation of VASP (Holt, Critchley and Brindle, 1998; Benz *et al.*, 2009), enabling its interaction with FA-associated proteins such as FAK, which have important roles in potentiating Rac1 activation and translocation to newly forming focal adhesions at the leading edges of cells (Kragtorp and Miller, 2006; Chang *et al.*, 2007). Taken together, these results suggest that NRP2’s modulatory role over Rac1 during focal adhesion assembly is dependent upon its ability to promote VASP-mediated actin remodelling.

### NRP2 promotes FAK phosphorylation and recruitment to assembling focal adhesion sites in ECs

We next considered whether NRP2 depletion would impair FAK phosphorylation. NRP2 has been demonstrated to localise to *p*-FAK-positive adhesions in breast-tumour epithelial cells allowed to adhere on laminin matrices (Goel *et al.*, 2012), and to regulate FAK signalling during branching morphogenesis in the developing mammary gland (Goel *et al.*, 2011). It is however unclear whether NRP2 influences FAK-mediated responses in microvascular ECs during angiogenesis. Following VEGF stimulation, FAK undergoes autophosphorylation at its Tyr^397^ residue, exposing binding sites for Src family kinases, which are involved in phosphorylating additional sites on FAK such as Tyr^407^ (Schaller, 2010; Herzog *et al.*, 2011). Both residues have been implicated in promoting EC migration and adhesion by potentiating the activation of downstream signalling cascades, such as the β-PAK interacting exchange factor (PIX)-dependent activation of Rac1 (Chang *et al.*, 2007). To assess whether NRP2 regulates FAK phosphorylation at either Tyr^397^ or Tyr^407^ in mLMECs adhered to FN, we assessed the expression of total and *p*-FAK in a similar manner as described above for VASP. At 5 minutes VEGF stimulation, NRP2 depleted ECs exhibited a less robust pattern of FAK phosphorylation at both Tyr^397^ and Tyr^407^ compared to Ctrl ECs (Suppl. Fig.1A-B, Fig.2A-B).

**Figure 2:**
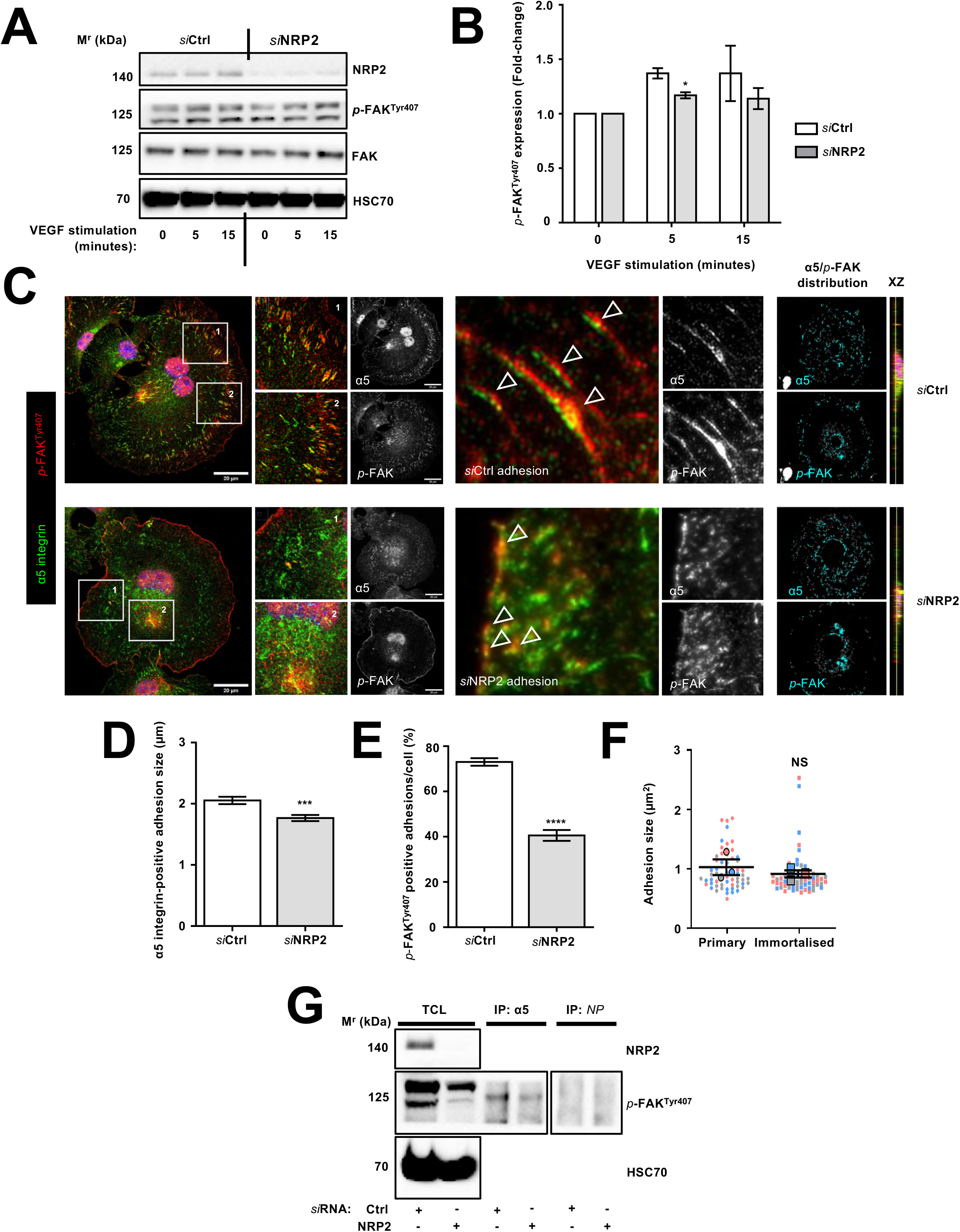
NRP2 promotes FAK phosphorylation and recruitment to assembling focal adhesion sites in ECs. **A**) siRNA-transfected ECs were seeded onto FN and prepared as described in **Fig.1D**. EC extracts were immunoblotted using antibodies against NRP2, *p*-FAK (Tyr407), FAK and HSC70. Panel shows representative western blot image. **B**) Densitometric quantification of mean *p*-FAK band intensities normalised against total FAK expression and obtained using ImageJ™. Error bars show ±SEM; N=3 independent experiments. Asterisks indicate statistical significance from an unpaired two-tailed *t*-test. **C**) siRNA-transfected ECs were seeded onto FN coated coverslips and allowed to adhere for 90 minutes at 37°C and 5% CO_2_ before being fixed. ECs were then co-immuno-stained for α5-integrin and *p*-FAK (Tyr407). Images were taken using a Zeiss LSM880 Airyscan Confocal microscope at 63X magnification. Arrowheads show co-localisation at peripheral adhesions. XZ plane images for each panel are also shown to display subcellular co-localisation. **D**) Quantification of mean α5-integrin positive adhesion size (microns) between Ctrl siRNA and NRP2 siRNA-treated ECs. Error bars show ±SEM; N=3 independent experiments; n≥180 adhesions/siRNA-treated group. Asterisks indicate statistical significance from an unpaired two-tailed *t*-test. **E**) Quantification of % of *p*-FAK (Tyr407)-positive α5-integrin adhesions per cell between Ctrl siRNA and NRP2 siRNA-treated ECs. Error bars show ±SEM; N=3 independent experiments; n≥30 cells/siRNA-treated group. Asterisks indicate statistical significance from an unpaired two-tailed *t*-test. **F**) Superplot showing quantification of mean adhesion area (microns^2^) per cell between primary and immortalised mLMECs. Error bars show ±SEM; N=3 independently derived EC clones (points from each biological replicate are shown as a different colour where the mean value for each replicate is shown as a larger circle of the same colour); n≥60 cells/ group. NS=not significant from an unpaired two-tailed *t*-test. **G**) siRNA-transfected ECs were seeded onto FN and allowed to adhere for 48 hours at 37°C and 5% CO_2_. EC extracts were immunoprecipitated by incubation with protein-G Dynabeads^®^ coupled to a α5-integrin antibody. Immunoprecipitated complexes were subjected to western blot analysis using antibodies against *p*-FAK (Tyr407). NRP2 silencing was confirmed by subjecting the total cell lysate to western blot analysis and incubating blots in antibodies against NRP2 and HSC70.

Tethering of α5-integrin to FN at assembling adhesion sites is required for initial FAK recruitment, aggregation and phosphorylation during nascent FA formation (Michael *et al.*, 2009; Schaller, 2010). We have previously shown that NRP2 interacts with and regulates the recycling of α5-integrin in ECs to promote the assembly of FAs at the leading edge (Alghamdi *et al.*, 2020). As NRP2 depletion impaired FAK phosphorylation, we sought to determine whether NRP2 also regulates the recruitment of FAK to assembling α5-integrin adhesions. As NRP1 disseminates VEGF-mediated signalling via the phosphorylation of FAK at Tyr^407^ (Herzog *et al.*, 2011), we focussed on how NRP2 regulates the interactions between α5-integrin and this residue specifically by immunofluorescence confocal microscopy. To this end, we examined the co-localisation between *p*-FAK^Tyr407^ and α5-integrin in Ctrl or NRP2-siRNA treated ECs allowed to adhere to FN for 90 minutes. Whilst Ctrl ECs exhibited a strong co-localisation between α5-integrin and *p*-FAK^Tyr407^ around the cell periphery within large characteristic adhesion structures, NRP2 depleted ECs displayed smaller α5-integrin adhesions, and those present were less enriched in *p*-FAK^Tyr407^. In addition, in NRP2 depleted cells we observed an increased accumulation of *p*-FAK^Tyr407^ around the perinuclear region (Fig.2C-E). To confirm that these differences did not arise as an artefact of cell immortalisation, we quantified the mean adhesion size in wild-type (WT)-derived primary and immortalised ECs adhered to FN for 90 minutes; we observed no significant alterations (Fig.2F). To corroborate these findings, we biochemically assessed the physical interaction between α5-integrin and *p*-FAK^Tyr407^ in Ctrl and NRP2 siRNA-treated ECs by co-immunoprecipitation. Lysate collected from Ctrl ECs displayed a more robust interaction between α5-integrin and *p*-FAK^Tyr407^ compared to lysate collected from NRP2 depleted ECs (Fig.2G), confirming that NRP2 is required for the proper recruitment and phosphorylation of FAK during FA assembly.

### NRP2 regulates an α5-integrin-*p*-FAKTyr407-Rab11 trafficking axis

Studies have revealed that the retention of *p*-FAK within endocytosing integrin complexes, including subsequent Rab11-associated recycling, sustains the active integrin conformation, enabling enhanced polarised reassembly of nascent FAs at the leading edge of the cell (Nader, Ezratty and Gundersen, 2016). We have previously provided evidence that NRP2 forms associations with Rab11 to facilitate α5-integrin recycling (Alghamdi *et al.*, 2020). As NRP2 depletion impaired the recruitment and phosphorylation of FAK to α5-integrin adhesion sites, we sought to determine whether NRP2 regulates an α5-integrin-*p*-FAK-Rab11 recycling axis to promote FA assembly and development. To examine this, we first stained for both Rab11 and NRP2 to ascertain their co-localisation in mLMECs. NRP2 colocalised with Rab11 at both the perinuclear region of the cell, and within trafficking vesicles (Suppl. Fig.2A), confirming our previous co-immunoprecipitation studies and suggesting a physical interaction.

To visualise the functional consequences of NRP2 silencing on Rab11-associated trafficking, we co-immuno-labelled Rab11 with endogenous *p*-FAK^Tyr407^ or α5-integrin in Ctrl or NRP2 siRNA-treated ECs allowed to adhere on FN for 90 minutes. As before, with NRP2 knockdown we observed a significant accumulation of *p*-FAK^Tyr407^ arresting within the perinuclear region of the cell, and a weaker co-localisation with fewer Rab11^+^ vesicles undergoing recycling back to assembling adhesion sites at the periphery of the cell (Fig.3A, C-E). Furthermore, NRP2 depleted ECs exhibited a dramatic un-coupling of α5-integrin with Rab11^+^ vesicles at both the perinuclear region and at the cell periphery (Fig.3B-C, F). These data suggest that NRP2 regulates Rab11-facilitated transport of both α5-integrin and *p*-FAK in ECs. Upon NRP2 silencing, assembling nascent adhesions are smaller, resulting from a reduced rate of α5-integrin recycling and *p*-FAK^Tyr407^ recruitment, concomitant with the accumulation of both α5-integrin and *p*-FAK^Tyr407^ at the perinuclear region.

**Figure 3:**
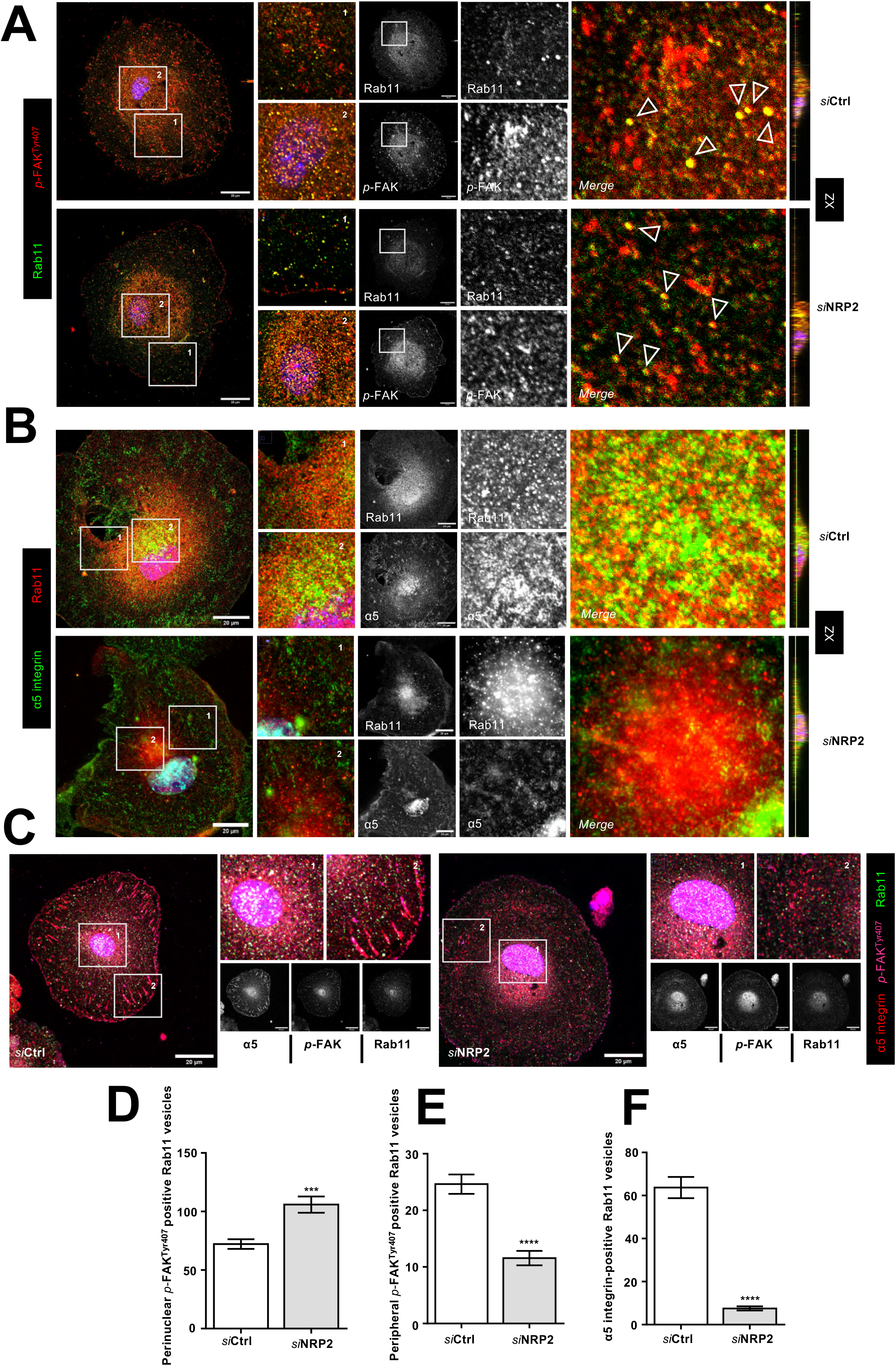
NRP2 regulates a α5-integrin-*p*-FAK^Tyr407^-Rab11 trafficking axis. **A**) siRNA-transfected ECs were seeded onto FN coated coverslips and allowed to adhere for 90 minutes at 37°C and 5% CO_2_ before being fixed. ECs were then co-immuno-stained for *p*-FAK (Tyr407) and Rab11. Images were taken using a Zeiss LSM880 Airyscan Confocal microscope at 63X magnification. Arrowheads show co-localisation. XZ plane images for each panel are also shown to display subcellular co-localisation. **B**) As described in **A**), ECs were co-immuno-stained for α5-integrin and Rab11. **C**) As described in **A**) ECs were co-immuno-stained for α5-integrin, *p*-FAK (Tyr407) and Rab11. **D**) Quantification of the mean number of perinuclear *p*-FAK (Tyr407)-positive Rab11 vesicles per 3x regions of interest (ROIs) between Ctrl siRNA and NRP2 siRNA-treated ECs. Error bars show ±SEM; n≥10 cells/siRNA-treated group. Asterisks indicate statistical significance from an unpaired two-tailed *t*-test. **E**) Quantification of the mean number of peripheral *p*-FAK (Tyr407)-positive Rab11 vesicles per 3x regions of interest (ROIs) between Ctrl siRNA and NRP2 siRNA-treated ECs. Error bars show ±SEM; n≥10 cells/siRNA-treated group. Asterisks indicate statistical significance from an unpaired two-tailed *t*-test. **F**) Quantification of the mean number of α5-integrin-positive Rab11 vesicles per cell between Ctrl siRNA and NRP2 siRNA-treated ECs. Error bars show ±SEM; N=3 independent experiments; n≥35 cells/siRNA-treated group. Asterisks indicate statistical significance from an unpaired two-tailed *t*-test.

### Long term depletion of NRP2 accelerates FA maturation and fibrillogenesis

As FAs mature and become influenced by actomyosin-driven forces, their composition transitions from small highly tyrosine phosphorylated focal contacts, to large macromolecular assemblies (Rossier *et al.*, 2012; Ajeian *et al.*, 2016). Whilst focal contacts often translocate by extending centripetally and contracting peripherally, and are rapidly turned over to promote migration, mature FA complexes anchor to the actin cytoskeleton to mediate more stable mechanical changes within the cell. Stress-fibre-associated FAs can further mature into fibrillar adhesions after approximately 24 hours, whereby engaged α5β1-integrin is translocated along actin cables centripetally towards the cell body. These elongated beaded structures become increasingly enriched with the actin-binding protein tensin-1 as they extend from the medial ends of the stationary FA, and act to funnel the necessary actomyosin tension required to unfold and integrate secreted FN dimers into an extracellular fibrillar network (Zamir *et al.*, 2000; Mana *et al.*, 2016). Fibrillar adhesion formation is therefore characteristic of a reduced motility and increased cell stability.

Since NRP2 promotes initial FA organisation and development by regulating Rab11-mediated trafficking of α5-integrin and FAK, we considered whether its protracted depletion would continue to impair FA maturation. Surprisingly, NRP2 siRNA-treated ECs adhered to FN for 180 minutes and 16 hours exhibited significantly larger α5-integrin positive adhesions than their Ctrl siRNA-treated counterparts (Fig.4A). Whilst Ctrl siRNA-treated ECs fixed at 16 hours exhibited mature, α5-integrin adhesions at the cell periphery, NRP2 depleted ECs displayed a high density of characteristic, tensin-1 positive fibrillar adhesions. Confocal XZ sectioning subsequently confirmed the co-localisation between endogenous α5-integrin and tensin-1 at the cell body in ECs depleted of NRP2, while in our Ctrl ECs tensin-1 exclusively co-localised with α5-integrin at peripheral punctae. This accelerated transition from mature FAs to fibrillar adhesions was found not to result from any changes in total tensin-1 expression at 16 hours, however NRP2 depleted ECs were found to express significantly more tensin-1 at 90 minutes (Fig.4B-D).

**Figure 4:**
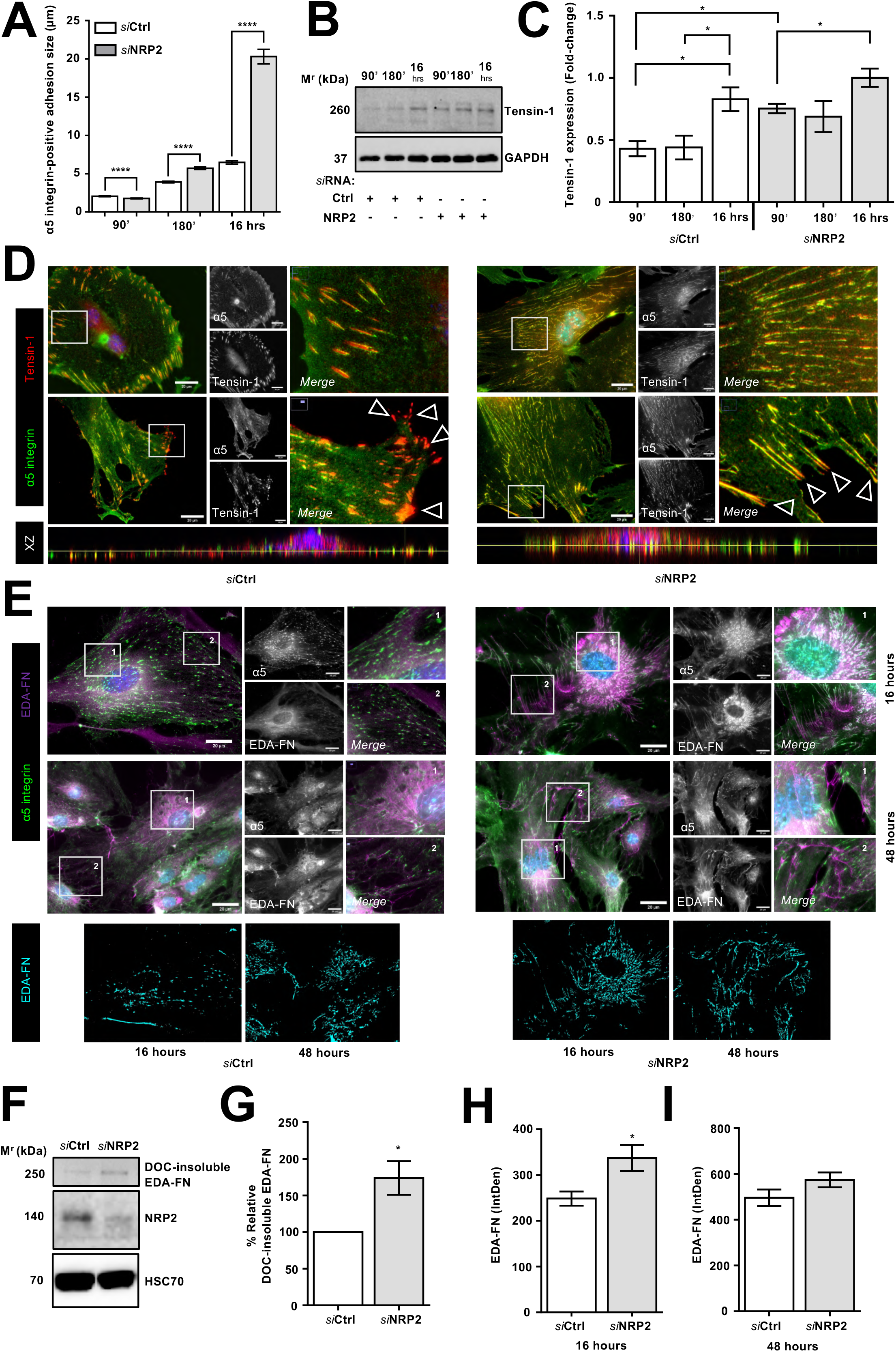
Long term depletion of NRP2 accelerates FA maturation and fibrillogenesis. **A)** Mean length of α5-integrin adhesions (μm), error bars show mean ± SEM; n≥150 adhesions/timepoint/condition); Asterisks indicate statistical significance from an unpaired two-tailed *t*-test. **B**) siRNA-transfected ECs were seeded onto 6 cm dishes pre-coated with 10 μg/ml FN and incubated for either 90 minutes, 180 minutes or 16 hours at 37°C. ECs were lysed in ESB and subjected to the DC protein assay before being analysed by Western blotting using primary antibodies against Tensin-1. GAPDH was used as a loading control. **C**) Bands were quantified using ImageJ™ densitometric analysis. Error bar shows mean ± SEM; N=3 independent experiments; asterisks indicate statistical significance from an unpaired two-tailed *t*-test. **D**) siRNA-transfected ECs were seeded onto FN coated coverslips and allowed to adhere for 16 hours at 37°C and 5% CO_2_ before being fixed. ECs were co-immuno-stained for α5-integrin and tensin-1. Images were taken using a Zeiss LSM880 Airyscan Confocal microscope at 63X magnification. XZ plane images for each panel are also shown to display colocalisation. **E**) As described in D), ECs were allowed to adhere for either 16 hours for 48 hours, before being co-immunolabelled with α5-integrin and EDA-FN. **F**) siRNA-treated ECs were allowed to adhere for 16 hours at 37°C and 5% CO_2._ ECs were then lysed, cleared and the insoluble fraction isolated. Insoluble fractions were separated by SDS-PAGE and subjected to Western blot analysis. Membranes were incubated in anti-EDA-FN primary antibody to assess quantities of insoluble cell secreted FN, anti-NRP2 primary antibody to confirm NRP2 depletion, and anti-HSC70 antibody as a loading control. **G**) % insoluble EDA-FN. Mean densitometric analysis obtained using ImageJ™. Error bars show mean ± SEM; N≥3 independent experiments; asterisks indicate statistical significance from an unpaired two-tailed *t*-test. **H/I**) Quantification of EDA-FN density at either 16 hours (**H**) or 48 hours (**I**) adhesion to FN. Error bars show mean ± SEM; n≥20 ECs/group. Asterisks indicate statistical significance from an unpaired two-tailed *t*-test.

Since an extended period of NRP2 depletion appeared to promote fibrillar adhesion formation, we asked whether this would also influence the polymerisation and incorporation of secreted FN into its ECM. We immunolabelled siRNA-treated ECs fixed at 16 hours with antibodies against α5-integrin and extra domain-A (EDA)-containing cellular FN (EDA FN), which has previously been employed to examine cell secreted FN specifically (Mana *et al.*, 2016; Sundararaman *et al.*, 2020). Confocal microscopy revealed that NRP2 silencing significantly increased EDA-FN secretion from the cell body and from the ends of fibrillar adhesions. We subsequently assessed the relative quantity of incorporated polymerised EDA-FN in lysates collected from both Ctrl and NRP2 siRNA treated ECs biochemically, and found that NRP2 depleted ECs exhibited significantly increased levels of insoluble EDA-FN (Fig.4E-G). We therefore infer that the initial aberrations in α5-integrin trafficking and FA turnover rate elicited by loss of NRP2 force FAs to instead mature at an accelerated rate into fibrillar adhesions. This is accompanied by a concomitant surge in FN fibrillogenesis.

It is worth noting, however, that when siRNA-treated ECs were allowed to adhere for 48 hours, we did not observe any differences in EDA-FN secretion or α5-integrin morphology (Fig.4E, H-I), suggesting that any aberration elicited from NRP2 silencing becomes compensated for by this timepoint.

### Loss of endothelial NRP2 inhibits tumour angiogenesis *in vivo*

Upregulations in FAK activity giving rise to aggressive tumour phenotypes has been well substantiated (Golubovskaya *et al.*, 2009; Jianliang Zhang and Steven N. Hochwald, 2014; Gao *et al.*, 2015; Lenzo and Cance, 2017). FA assembly and subsequent FAK phosphorylation events in turn play a crucial role in FA turnover and remodelling of the actin cytoskeleton to promote cancer cell metastasis (Nagano *et al.*, 2012). NRP2 overexpression is also known to promote tumorigenicity and metastasis in a multitude of cancer subtypes. It therefore represents a potential diagnostic or prognostic biomarker and therapeutic target for inhibiting primary tumour growth (Borkowetz *et al.*, 2020). Despite this, little investigation into its contribution during tumour vascularisation has been made. To isolate the role of NRP2 during tumour angiogenesis, we crossed NRP2-floxed (NRP2^*flfl*^)-mice to tamoxifen (OHT)-inducible PDGFb-iCreERT2 mice and examined the effect of an acute EC-specific depletion of NRP2 on subcutaneous allograft tumour growth using CMT19T lung carcinoma cells. Cells were allowed to grow for a period of 18 days alongside thrice weekly injections of tamoxifen to effectively deplete NRP2 expression within the tumour vasculature (Fig.5A), avoiding the complications of interpreting NRP2 function using global knockout models. By tracking changes in tumour volume from day 10 post implantation, it was also possible to examine any temporal limitations to depleting NRP2 in this manner.

**Figure 5:**
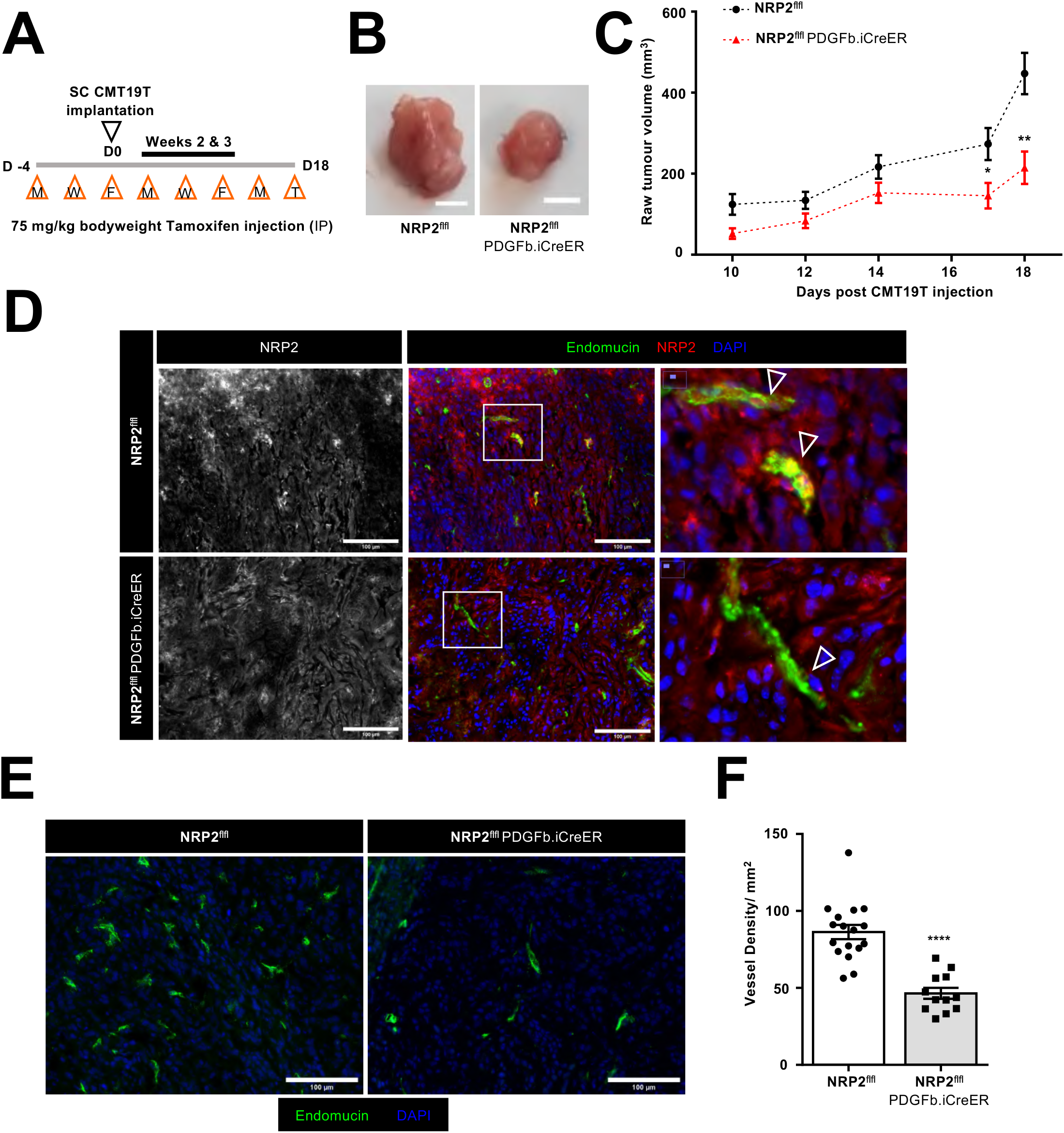
Loss of endothelial NRP2 inhibits tumour angiogenesis *in vivo*. Inducible, endothelial specific deletion of NRP2 was achieved by crossing mice expressing the PDGFb.iCreER promoter of Cre-recombinase to those floxed for NRP2. **A**) Experimental schematic: tamoxifen-induced activation of Cre-recombinase and thus deletion of NRP2 was employed via the following regimen in order to study its role during tumorigenesis. NRP2^*flfl*^; PDGFb.iCreER (Cre-positive) and NRP2^*flfl*^ (Cre-negative) littermate control mice received intraperitoneal (IP) injections of tamoxifen (75 mg/kg bodyweight) thrice weekly (Monday, Wednesday, Friday) for the duration of the experiment from D-4 to D17 to induce Cre-recombinase activity. CMT19T lung carcinoma cells (1×106) were implanted subcutaneously (SC) into the flank of mice at D0 and allowed to grow until D18. **B)** Representative images of tumours harvested from Cre-negative and Cre-positive mice at day 18. **C**) Quantification of tumour volumes measured from tumour bearing Cre-negative and Cre-positive mice measured between day 10 and day 18 post CMT19T injection. Error bars show mean ± SEM; N=3 independent experiments; n≥12 tumours per genotype. Asterisks indicate statistical significance from an unpaired two-tailed *t*-test. **D**) IHC staining using Endomucin and NRP2 primary antibodies to confirm endothelial specific NRP2 deletion in blood vessels from Cre-positive tumours. **E**) Representative tumour sections from Cre-negative and Cre-positive tumours showing endomucin-positive blood vessels. **F**) Quantification of number of blood vessel density per mm^2^. Mean quantification performed on 3x ROIs per tumour section, from 3x sections per tumour. Error bars show mean ± SEM; N=3 independent experiments; n≥12. Asterisks indicate statistical significance from an unpaired two-tailed *t*-test.

Tumours grown in NRP2^flfl^.PDGFb-iCreERT2-positive (Cre-positive) animals developed significantly smaller than their control littermate NRP2^flfl^ (Cre-negative) counterparts despite no gross changes in mean animal weight (Fig.5B-C, Suppl. Fig3A). To directly assess the effects of EC-depletion of NRP2 on tumour angiogenesis, we performed immuno-fluorescent analysis on all tumours harvested from Cre-negative and Cre-positive mice. After confirming endothelial specific depletion of NRP2 within endomucin-positive blood vessels in Cre-positive tumour sections (Fig.5D), we found that compared to tumours from Cre-negative animals, those deficient in endothelial NRP2 displayed significantly less vasculature (Fig.5E-F). This indicates that the supressed tumour growth and reduced tumour angiogenesis we observe in Cre-positive mice likely results from a NRP2-specific, EC-intrinsic defect.

### Deficiency of endothelial NRP2 transiently impairs neonatal retinal vascular development

Whilst pathological and physiological angiogenesis are both driven by many of the same processes and stimuli, tumour vasculature is very much distinct from physiological vasculature. In addition to their highly tortuous organisation, vessels that form during tumorigenesis are far leakier than those that form under physiological homeostasis (Deshpande, Biswas and Torchilin, 2013; Ghosh Dastidar, Ghosh and Chakrabarti, 2020). Studying the effects of depleting endothelial NRP2 in a physiological model was therefore necessary to obtain any mechanistic insight into its function during developmental angiogenesis. To this end, we utilised the postnatal mouse retina to examine whether any defect arose from inducing EC-specific depletion of NRP2 at postnatal day 6 (P6) or P12, timepoints during which the superficial plexus and the deep plexus are forming respectively (Milde *et al.*, 2013). Following tamoxifen administration regimens, we first confirmed successful depletion of NRP2 by co-staining with the endothelial marker BS-1 lectin in retinas harvested from Cre-negative and Cre-positive mice (Fig.6A).

**Figure 6:**
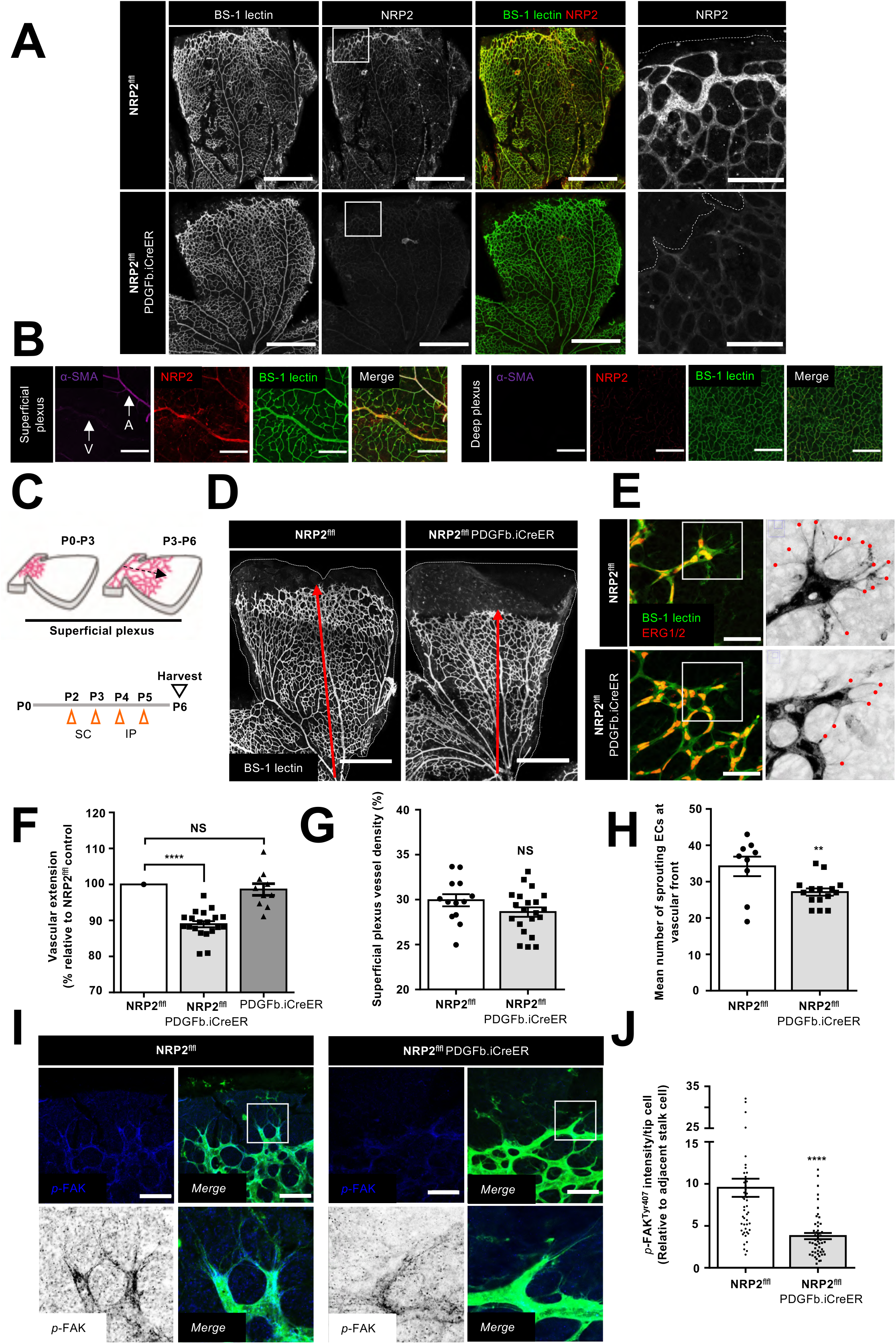
Deficiency of endothelial NRP2 transiently impairs neonatal retinal vascular development. Inducible, endothelial specific deletion of NRP2 was achieved by crossing mice expressing the PDGFb.iCreER promoter of Cre-recombinase to those floxed for NRP2. **A**) Tamoxifen-induced NRP2 deletion was confirmed by co-staining NRP2 with BS1-lectin. Scale bar = 500 μm. **B**) Representative images of BS-1 lectin, α-SMA and NRP2 labelled vasculature from Cre-negative and Cre-positive retinas harvested at P6 and P12. Panels show superficial and deep plexus layers. Scale bar = 150 μm. **C**) Experimental schematic: tamoxifen-induced activation of Cre-recombinase and thus deletion of NRP2 was employed via the following regime in order to study its role during the formation of the developing retinal superficial vascular plexus. Cre-positive and Cre-negative littermate control mice received subcutaneous (SC) injections of tamoxifen (50 μl, 2mg/ml stock) on postnatal (P) days 2 and 3, followed by intraperitoneal (IP) injections of the same dose on P4 and 5. Retinas were then harvested at P6 (Figure adapted from Milde *et al.*, 2013). **D**) Representative images of BS-1 lectin labelled P6 retinas harvested from Cre-negative and Cre-positive mice showing vascular extension from the optic nerve. Scale bar = 500 μm. **E**) Representative images of BS-1 lectin and ERG1/2 co-labelled P6 retinas showing sprouting ECs at the vascular front. Scale bar = 50 μm. Grayscale images show magnified ROIs from left panels, with filopodial extensions highlighted in red. **F**) Quantification of P6 vascular extension from the optic nerve, presented as a percentage of the average extension observed in their Cre-negative littermate controls. Error bars show mean ± SEM; N=3 independent experiments; n≥10 retinas. Asterisks indicate statistical significance from an unpaired two-tailed *t*-test. **G**) Quantification of mean % vessel density from 350×350 μm ROIs taken per leaf of each P6 retina. Error bars show mean ± SEM; N=3 independent experiments; n≥13 retinas. NS stands for non-significance from an unpaired two-tailed *t*-test. **H**) Quantification of the mean number of sprouting ECs per retina leaf. Error bars show mean ± SEM; N=2 independent experiments; n≥10 retinas. Asterisks indicate statistical significance from an unpaired two-tailed *t*-test. **I**) Representative images of *p*-FAK Tyr-407 labelled retinal vasculature co-labelled with BS-1 lectin from Cre-negative and Cre-positive mice at P6. Scale bar = 50 μm. **J**) Quantification of *p*-FAK Tyr-407 intensity within sprouting tip cells made relative to adjacent stalk cell intensities. Error bars show mean ± SEM; n≥50 tip cells); ****=P<0.0001, unpaired students t-test (two-tailed).

The expression of NRP2 is largely considered to be more venous than arterial within the endothelium (Bielenberg *et al.*, 2006; Fantin *et al.*, 2011). We were therefore unsurprised to observe enriched NRP2 expression within veins of the superficial plexus by co-immuno-labelling with α-smooth muscle actin (α-SMA), a marker of smooth muscle cells that are known to primarily ensheathe arteries (Alarcon-Martinez *et al.*, 2018). We also detected NRP2 expression in the disseminating microvasculature of the superficial plexus, particularly at the vascular front. Subsequent multi-plexus analysis revealed, however, very little NRP2 expression in the deep vascular layer at P12, suggesting that NRP2 plays a more prominent role during initial vessel sprouting outwards from the optic nerve to the retinal periphery rather than during deep plexus development (Fig.6A-B). To examine this further, we compared the vascular extension from the optic nerve between retinas harvested from Cre-negative and Cre-positive mice and found that EC-specific depletion of NRP2 significantly reduced radial expansion of the vasculature at P6 without affecting vessel density. By comparing the vascular extension of Cre-negative and PDGFb-iCreERT2-positive (Cre-only) retinas harvested at P6, and observing no significant differences (Suppl. Fig.4A), we also confirmed that this phenotype did not arise due to Cre-toxicity (Brash *et al.*, 2020). Cre-positive retinas were also found to exhibit fewer sprouting ECs at the vascular front, consistent with NRP2’s expression profile in Cre-negative retinas within the superficial layer (Fig.6C-H). Co-staining between BS-1 lectin and collagen IV also revealed that Cre-positive retinas exhibited significantly more vessels undergoing regression within the superficial plexus. This, paired with a significant reduction in vessel diameter, suggests that the vasculature of Cre-positive mice (Suppl. Fig.4B-D) is less functional than that of their Cre-negative counterparts.

Whilst the existence of FA structures in the developing postnatal retina is disputed, it has been demonstrated that FA-associated signalling cascades do exist (Raimondi *et al.*, 2014; Schimmel *et al.*, 2020). In an attempt to corroborate our *in vitro* findings, we sought to examine differential FAK phosphorylation in the retina of our NRP2 deficient mice. Confocal microscopy of retinas harvested at P6 revealed that *p*-FAK^Tyr407^ staining was greatest at the vascular front and enriched at the apical ends of sprouting tip cells. *p*-FAK^Tyr407^ was also detected along the length of extending filopodia. However, in retinas harvested from Cre-positive mice, the relative intensity of *p*-FAK^Tyr407^ staining in sprouting tip cells was significantly diminished (Fig.6I-J). Taken together, this suggests that the reduced vascular extension and increased vascular regression we observe following NRP2 depletion, likely results from the inability of NRP2-deficient ECs to form stable cell-matrix interactions.

Defects at P6 were found to be transient however, as at P12, retinas from both Cre-negative and Cre-positive mice exhibited fully vascularised superficial plexus layers, with no gross differences in vessel density (Suppl. Fig.4E-H). As we detected far less robust levels of NRP2 expression in the deep plexus layer of Cre-negative retinas compared to at the superficial plexus, we were also unsurprised to see the development of a normal vascular bed at the deep plexus layer in Cre-positive mice (Suppl. Fig.4I-J). These data suggest that NRP2 is primarily involved in promoting initial EC sprouting and vascularisation of the superficial plexus rather than during deep plexus development.

## Discussion

When considered in conjunction with their interactions with other receptors known to regulate angiogenesis, such as α5β1- and αvβ3-integrins, the evidence purporting both NRPs as pro-angiogenic molecules holds true. For example, Valdembri et al., have previously reported that NRP1 promotes α5β1-integrin-mediated adhesion to FN matrices by selectively stimulating its rapid endocytosis from fibrillar adhesions and subsequent Rab5-dependent recycling to drive FA turnover in ECs (Valdembri *et al.*, 2009). Similarly, both we and others have demonstrated that NRP2 promotes cellular adhesion and migration via its interactions with the α5-integrin subunit (Cao *et al.*, 2013; Alghamdi *et al.*, 2020). Further evidence for this has recently been provided by Luo et al., who demonstrated that NRP2 may regulate migration by upregulating cofilin activity, a key mediator of actin depolymerisation at slow growing filament ends (Luo *et al.*, 2020). Here, we provide evidence that NRP2 associates with a number of other proteins known to stabilise and promote actin branch nucleation and stress fibre growth, notably cortactin, VASP and fascin (Woo and Fowler, 1994; Yamakita *et al.*, 1996; Uruno *et al.*, 2001; Benz *et al.*, 2009). It is likely therefore, that NRP2 drives the simultaneous depolymerisation of F-actin at slow-growing filament ends to supply the demand for new actin monomers at assembling fast growing ends of extending filopodial protrusions. NRP2-mediated VASP phosphorylation in turn potentiates the rapid polymerisation and nucleation of F-actin at fast growing ends by initiating the translocation and activation of Rac1 to newly forming adhesions.

It is known that aberrations in VASP activity result in reduced FAK phosphorylation. For example, VASP inhibition has been shown to significantly reduce phosphorylation at Tyr-925 in a human chronic myeloid leukaemia (CML) cell line, and decrease Tyr^397^ phosphorylation during Xenopus somite development (Kragtorp and Miller, 2006; Bernusso *et al.*, 2015). As VASP is strongly Ser^157^ phosphorylated at FAs (Holt, Critchley and Brindle, 1998; Benz *et al.*, 2009), we were unsurprised to observe a corresponding significant reduction in FAK phosphorylation at Tyr^397^ and Tyr^407^ sites, both implicated in canonical VEGF gradient-driven EC adhesion and migration through their necessity for correct assembly of nascent FAs. VEGF-mediated phosphorylation of FAK at its Tyr^407^ residue during integrin-directed adhesion and migration is also known to be NRP1 dependent (Chang *et al.*, 2007; Herzog *et al.*, 2011).

While NRP2 has been shown to direct FAK-mediated adhesion in cancerous epithelial cells via α6β1 (Goel *et al.*, 2012), and to support branching morphogenesis in the developing mammary gland (Goel *et al.*, 2011), no connection has been explored in microvascular ECs. Endogenous *p*-FAK^Tyr407^ staining in fixed ECs subsequently revealed that significantly less phosphorylated FAK appears to be recruited to α5-integrin positive adhesion sites following NRP2 knockdown, resulting in smaller less developed FAs. Given we have previously provided evidence to suggest that NRP2 exerts a mainly VEGF-independent role during EC adhesion and migration (Alghamdi *et al.*, 2020), we propose that NRP2 instead promotes FAK auto-phosphorylation following integrin engagement and clustering.

As NRP2 silencing disrupted Rab11-directed recycling of α5-integrin-*p*-FAK complexes back to assembling nascent adhesions, it is likely that NRP2 potentiates FAK-mediated signalling cascades by regulating the pool of available α5β1-integrin capable of binding the FN matrix. In support of this, studies have purported that α5-integrin is maintained in an active, unliganded conformation when endocytosed with active FAK, and during Rab11-mediated recycling (Nader, Ezratty and Gundersen, 2016). This then enables more efficient integrin engagement with the matrix, and subsequent reassembly of polarised FAs to promote directional migration. As we observed a significant loss in co-localisation between *p*-FAK^Tyr407^ and α5-integrin within Rab11^+^ vesicles, concomitant with an accumulation of *p*-FAK^Tyr407^ at the perinuclear region, we infer that NRP2 is involved in maintaining the interactions between active FAK and α5-integrin during vesicular transport.

To our surprise, protracted siRNA-mediated depletion of NRP2 accelerated the transition from mature FAs to tensin-1 positive fibrillar adhesions translocating α5-integrin to the cell body. We believe this to be an artefact arising from aberrations in FA turnover rate, whereby FAs are forced to mature prematurely rather than be dynamically endocytosed and recycled. To confirm this model, we examined the effects of NRP2 silencing on the assembly of fibrillar FN. This remodelling of the extracellular matrix is driven by the actomyosin tension accrued from the translocation of α5β1 integrin into fibrillar adhesions and is essential during angiogenesis for vessel formation (Mana *et al.*, 2016). Contemporaneous to the surge in fibrillar adhesion formation we observed in our NRP2 depleted ECs fixed at 16 hours, immunostaining and deoxycholate (DOC)-buffer extracted insoluble EDA-FN quantification revealed a surge in secreted FN radiating outwards from α5-integrin containing adhesions. In support of these findings, others have established that Rab11-mediated recycling of α5-integrin orchestrates the continuous renewal of polarised FN fibrils being secreted to form an extracellular fibrillar network (Mana *et al.*, 2016; Sundararaman *et al.*, 2020). We propose here that the early defects in FA turnover rate elicited by NRP2 depletion, although eventually compensated for, sustains the transition to a more sedentary cell fate in which the premature development of fibrillar adhesions promotes the integration of secreted FN into insoluble fibrils. By doing so, stability and contractility is favoured over motility. Alongside other studies (Bielenberg *et al.*, 2012), this work reinforces the growing evidence in multiple cell types that NRP2 deficiency can promote cell contractility.

In an effort to emphasise the relevance of endothelial NRP2 during angiogenesis and its coordination of α5-integrin trafficking *in vivo*, we modelled the effects of its EC specific depletion during both pathological and physiological vessel growth. Whilst it has been substantiated by many that NRP2 promotes a more aggressive cancer phenotype (Borkowetz *et al.*, 2020), our findings clearly demonstrate that the expansion of tumour vasculature to support primary tumour growth is inhibited following loss of endothelial NRP2. Likewise, in the developing retina, vascular outgrowth from the optic nerve at P6 is impaired in animals depleted in endothelial NRP2. NRP2 is largely considered to be expressed to a greater extent within veins as opposed to arteries (Bielenberg *et al.*, 2006), however little is known about its expression profile or its endothelial-specific contributions during the hierarchical vascularisation of the postnatal retina. Studies have shown that NRP2 does not compensate for the loss of NRP1’s cytoplasmic domain during retina development and arterial/venous differentiation (Fantin *et al.*, 2011). Indeed, the published evidence outlining the roles of the neuropilins during angiogenesis support the principle that NRP1 likely plays a more dominant role over NRP2. Despite this, we found that vascular extension within the superficial plexus was significantly impaired following NRP2 depletion at P6, owing to a decrease in the number of sprouting tip cells at the vascular front. These defects were found to likely arise from a deficiency in polarised FA-associated signalling, specifically FAK phosphorylation within sprouting tip cells, which has been previously alluded to be responsible for the formation of stable cell-matrix interactions (Schimmel *et al.*, 2020). These findings align with the expression profile of NRP2, which showed a far more robust expression at the vascular front of the superficial plexus, compared to within the deep plexus layer.

Taken together, these data presented provides evidence that endothelial NRP2 plays a key role in regulating both pathological and physiological angiogenesis. We propose a model whereby NRP2 promotes FA development by regulating α5-integrin trafficking and show that NRP2 is essential for vessel development by enabling the formation of stable cell-matrix connections.

## Supplementary Figure Legends

**Suppl. Fig.1.**
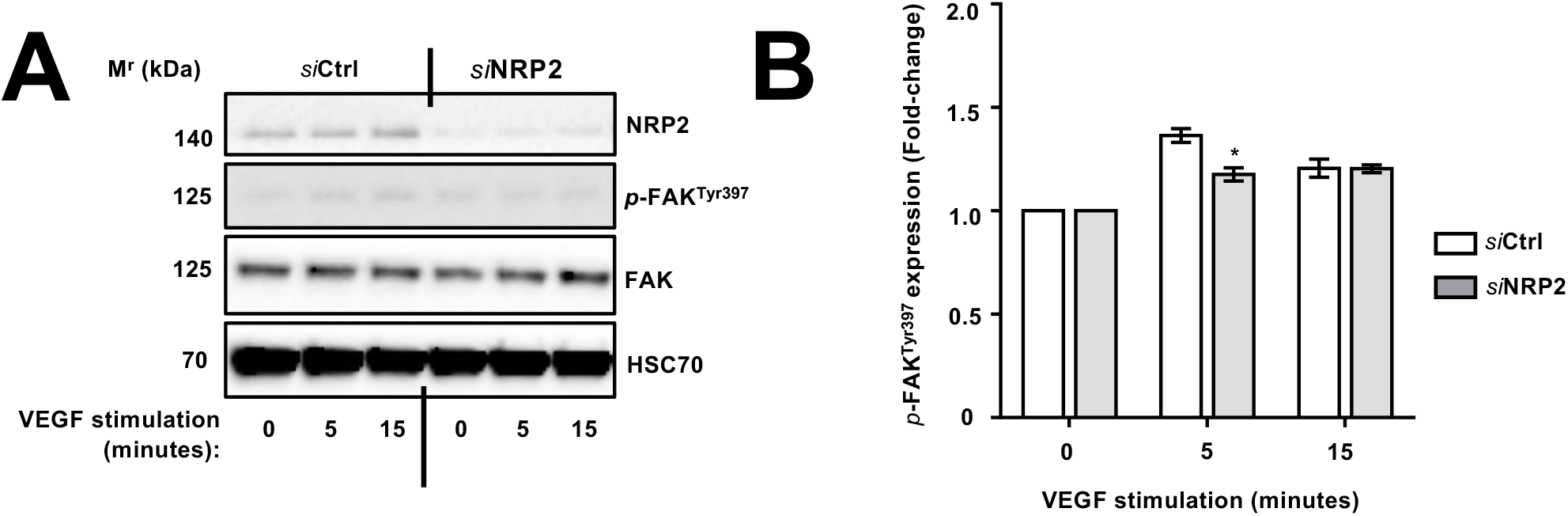
**A)** siRNA-transfected ECs were seeded onto FN and prepared as described in **Fig.1D**. EC extracts were immunoblotted using antibodies against NRP2, *p*-FAK (Tyr397), FAK and HSC70. Panel shows representative western blot image. **B**) Densitometric quantification of mean *p*-FAK band intensities normalised against total FAK expression and obtained using ImageJ™. Error bars show ±SEM; N=3 independent experiments. Asterisks indicate statistical significance from an unpaired two-tailed *t*-test.

**Suppl. Fig.2.**
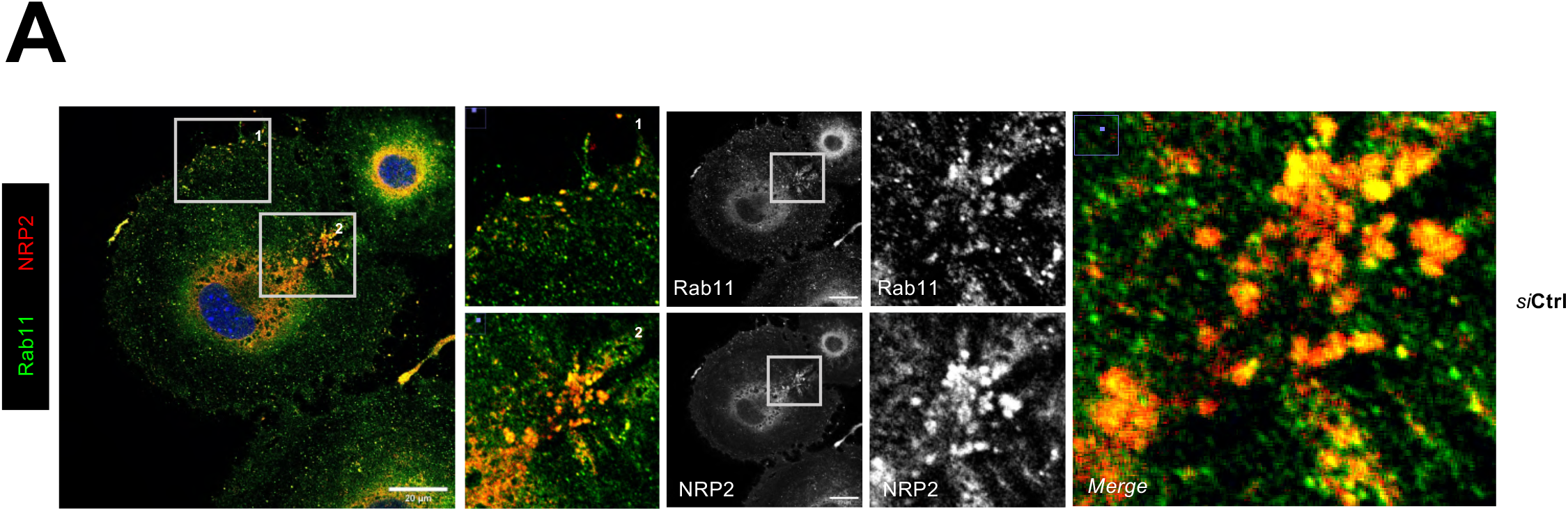
**A**) Ctrl siRNA-transfected ECs were seeded onto FN coated coverslips and allowed to adhere for 90 minutes at 37°C and 5% CO_2_ before being fixed. ECs were then co-immuno-stained for NRP2 and Rab11. Images were taken using a Zeiss LSM880 Airyscan Confocal microscope at 63X magnification. Panels show colocalisation at the perinuclear region and in trafficking vesicles at the periphery of cells.

**Suppl. Fig.3.**
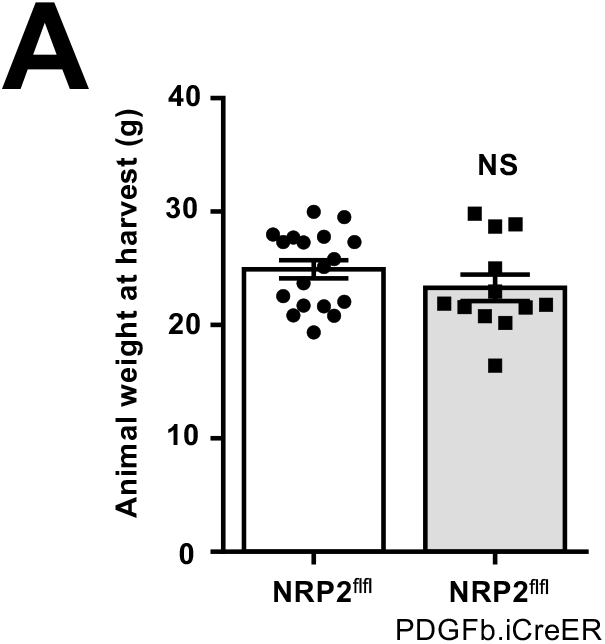
**A)** Quantification of mean animal weight (g) on day of harvest on day 18 post CMT19T injection. Error bars show mean ± SEM; N=3 independent experiments; n≥12.

**Suppl. Fig.4.**
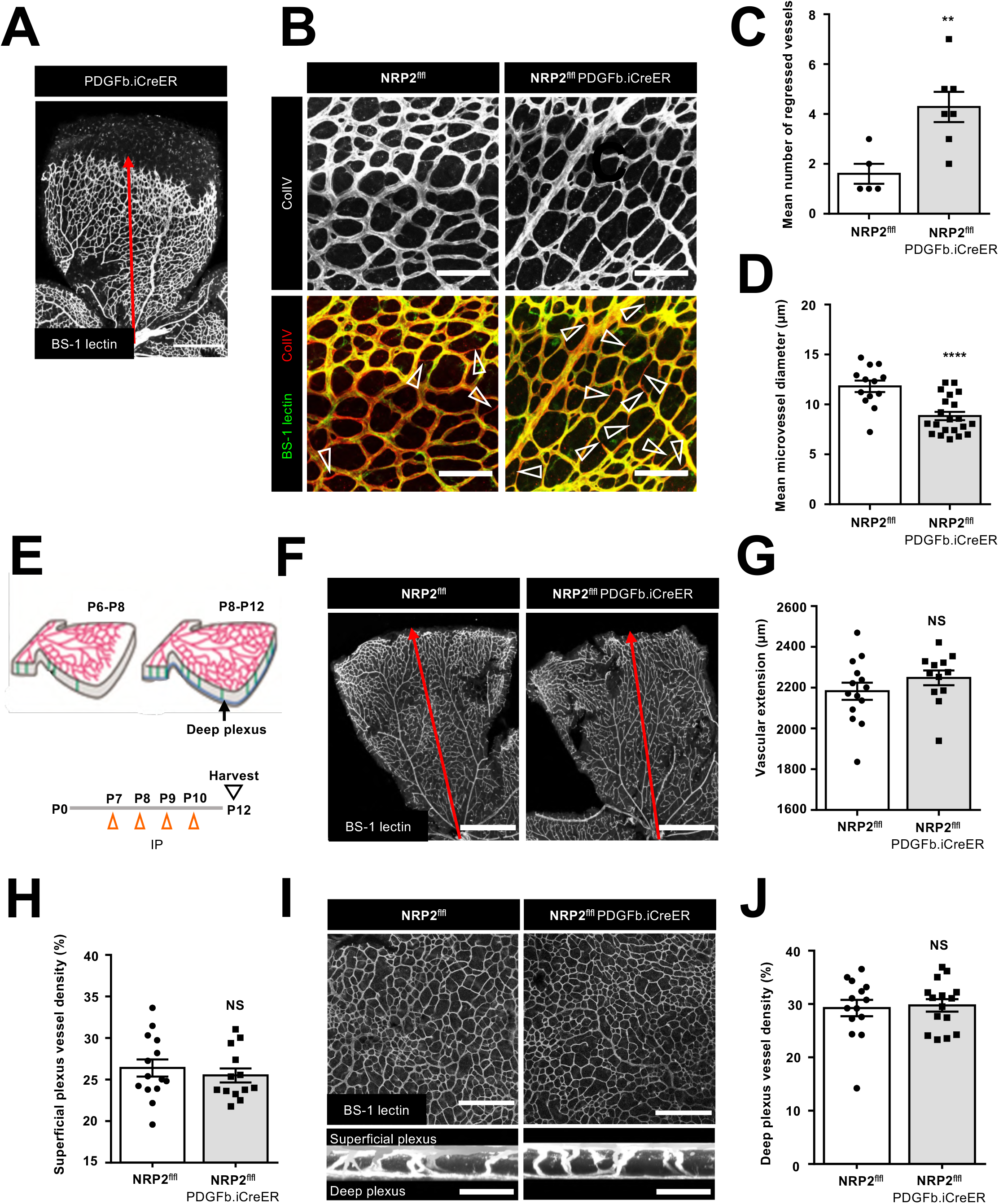
**A)** Representative image of a BS-1 lectin labelled P6 retina harvested from a Cre-only animal showing vascular extension from the optic nerve. Scale bar = 500 μm. **B)** Representative images of Collagen IV (ColIV) labelled vessel sleeves from P6 retinas harvested from Cre-negative and Cre-positive mice. Bottom panels show overlays with BS-1 lectin to identify ColIV positive BS-1 lectin negative sleeves. Arrowheads show regressed vessels. Scale bar = 150 μm. **C**) Quantification of the mean number of regressed vessels observed in 3× 200×200 μm ROIs taken per leaf of each retina. Error bars show mean ± SEM; N=2 independent experiments; n≥5 retinas. Asterisks indicate statistical significance from an unpaired two-tailed *t*-test. **D**) Quantification of mean micro-vessel diameter calculated from 350×350 μm ROIs taken per leaf of each retina. The mean diameter of 5 randomly selected vessels was measured per ROI. Error bars show mean ± SEM; N=2 independent experiments (n≥13 retinas). Asterisks indicate statistical significance from an unpaired two-tailed *t*-test. **E**) Experimental schematic: tamoxifen-induced activation of Cre-recombinase and thus deletion of NRP2 was employed via the following regimen in order to study its role during the formation of the developing retinal deep plexus. Cre-positive and Cre-negative littermate control mice received IP injections of tamoxifen (50 μl, 2mg/ml stock) on postnatal (P) days 7 to 10. Retinas were then harvested at P12 (Figure adapted from Milde *et al.*, 2013). **F**) Representative images of BS-1 lectin labelled P12 retinas harvested from Cre-negative and Cre-positive mice showing full vascular extension from the optic nerve. **G**) Quantification of P12 vascular extension from the optic nerve. Error bars show mean ± SEM; N=3; n≥12 retinas. NS stands for non-significance from an unpaired two-tailed *t*-test. **H**) Quantification of mean % vessel density from 350×350 μm ROIs taken per leaf of each P12 retina. Error bars show mean ± SEM; N=3 independent experiments; n≥14 retinas. NS = not significant from an unpaired two-tailed *t*-test. **I**) Top panels: representative images of BS-1 lectin labelled P12 retinas showing the deep plexus layer. Scale bar = 500 μm. Bottom panels: representative spliced Z stack images of BS-1 lectin labelled P12 retinas showing normal descending vessel sprouting and deep plexus layer formation. **J**) Quantification of mean % vessel density from 850×850 μm ROIs taken per leaf of each P12 retina. Error bars show mean ± SEM; N=3 independent experiments; n≥14 retinas. NS = not significant from an unpaired two-tailed *t*-test.

## Materials and Methods

### Animal Generation

All experiments were performed in accordance with UK home office regulations and the European Legal Framework for the Protection of Animals used for Scientific Purposes (European Directive 86/609/EEC), prior to the start of this project. NRP2 floxed (NRP2^flfl^) mice were purchased from Jackson Laboratories (Bar Harbour, Maine, USA), and were generated by gene target insertion of embryonic stem cells, resulting in the insertion of loxP sites flanking exon 1 of the NRP2 gene. A loxP-tauGFP FRT-flanked neo cassette was inserted 1 kb into intron 1 via homologous recombination. Heterozygous animals were crossed to an flp recombinase transgenic line for removal of the neo cassette. The PCR analysis to confirm floxing was carried out using the following oligonucleotide primers:

Forward (WT) primer (Reaction A): 5’-CAGGTGACTGGGGATAGGGTA -3’

Common primer (Reaction A + B): 5’-AGCTTTTGCCTCAGGACCCA -3’

Forward primer (^*flfl*^) (Reaction B): 5’-CCTGACTACTCCCAGTCATAG -3’

Transgenic mice expressing a tamoxifen-inducible PDGFb-iCreER^T2^ allele in vascular ECs were provided by Marcus Fruttiger (UCL, London, UK), and were generated by substituting the exon 1 of the PDGFb gene by the iCreER^T2^-IRES-EGFP-pA sequence.

Forward primer: 5’-GCCGCCGGGATCACTCTC-3’

Reverse primer: 5’-CCAGCCGCCGTCGCAACT-3’

NRP2^flfl^ mice were bred with PDGFb.iCreER^T2^ mice to generate NRP2^*flfl*^; PDGFb.iCreER animals. PDGFβ-iCreERT2 expression was maintained exclusively on breeding males, thereby ensuring the generation of both Cre-negative and positive offspring and enabling the use of littermate controls. All animals were bred on a pure C57/BL6 background.

### Cell Isolation, Immortalisation and Cell Culture

Primary mouse lung microvascular endothelial cells (mLMECs) were isolated from adult mice bred on a pure C57/BL6 background. Primary ECs were twice positively selected for through their expression of endomucin by magnetic activated cell sorting (MACS) as previously described by Reynolds & Hodivala-Dilke (Reynolds and Hodivala-Dilke, 2006). ECs were immortalised using polyoma-middle-T-antigen (PyMT) retroviral transfection as previously described by Robinson et al (Robinson *et al.*, 2009). Immortalised mLMECs were cultured in IMMLEC media, a 1:1 mix of Ham’s F-12:DMEM medium (low glucose) supplemented with 10% FBS, 100 units/mL penicillin/streptomycin (P/S), 2 mM glutamax, 50 μg/mL heparin (Sigma).

Immortalised mLMECs were cultured on 0.1% gelatin coated flasks at 37°C in a humidified incubator with 5% CO_2._ For experimental analyses, plates, dishes and flasks were coated in 10 μg/ml human plasma fibronectin (FN) (Millipore) overnight at 4°C. Vascular endothelial growth factor-A (VEGF-A_164_: mouse equivalent of VEGF-A_165_) was made in-house as previously described by Krilleke et al. (Krilleke *et al.*, 2007).

### siRNA Transfection

ECs were transfected with non-targeting control siRNA or a mouse-specific NRP2 siRNA construct (Dharmacon), suspended in nucleofection buffer (200 mM Hepes, 137 mM NaCl, 5 mM KCl, 6 mM D-glucose, and 7 mM Na_2_HPO_4_ in nuclease-free water; filter sterilised) using the Amaxa 4D-nucleofector system (Lonza) under nucleofection program EO-100 according to manufacturer’s instructions.

### Immunocytochemistry

siRNA-transfected ECs were seeded onto FN-coated acid-washed, oven sterilised glass coverslips in 24-well plates at a seeding density of 2.5×10^4^ cells/well. ECs were fixed at indicated timepoints in 4% paraformaldehyde (4% PFA) for 10 minutes, washed in PBS, blocked and permeabilised with 10% goat serum, PBS 0.3% triton X-100 for 1 hour at room temperature. Cells were incubated in primary antibody in PBS overnight at 4°C. Primary antibodies were: anti-α5-integrin (clone ab150361; Abcam), anti-p-FAK^Tyr407^ (clone 44-650G Invitrogen), anti-Rab11 (clone 3612; Abcam), anti-tensin-1 (clone NBP1-84129; Novus Biologicals), EDA-FN (F6140; Sigma). Coverslips were PBS washed, and incubated with an appropriate Alexa fluor secondary antibody diluted 1:200 in PBS for 2 hours at room temperature. F-actin staining was performed by incubating cells in phalloidin-364 diluted 1:40 in PBS for 2 hours at room temperature during secondary antibody incubation. Coverslips were PBS washed again, before being mounted onto slides with Prolong^^®^^ Gold containing DAPI (Invitrogen). Images were captured either using a Zeiss AxioImager M2 microscope (AxioCam MRm camera) at 63x magnification or using a Zeiss LSM880 Airyscan Confocal microscope at 63X magnification. FA number and size, and EDA-FN fibers/cell were quantified using ImageJ™ software as previously described by Lambert et al (Lambert *et al.*, 2020). XZ confocal sections taken across the z-plane were processed to form a 2D projection representing the full depth of the cell culture.

### Western Blot Analysis

siRNA transfected ECs were seeded into FN-coated 6-well plates at a seeding density of 5×10^5^ cells/well and incubated for 48 hours at 37°C in a 5% CO_2_ incubator. ECs were lysed in electrophoresis sample buffer (ESB) (Tris-HCL: 65 mM pH 7.4, sucrose: 60 mM, 3% SDS), and homogenised using a Tissue Lyser (Qiagen) with acid-washed glass beads (Sigma). Following protein quantification using the DC BioRad assay, 30 μg of protein from each sample was loaded onto 8% polyacrylamide gels and subjected to SDS-PAGE. Proteins were transferred to a nitrocellulose membrane (Sigma) and incubated in 5% milk powder in PBS 0.1% Tween-20 (0.1% PBST) for 1 hour at room temperature followed by an overnight incubation in primary antibody diluted 1:1000 in 5% bovine serum albumin (BSA) in 0.1% PBST at 4°C. Membranes were washed 3x with 0.1% PBST and incubated in an appropriate horseradish peroxidase (HRP)-conjugated secondary antibody (Dako) diluted 1:2000 in 5% milk powder in 0.1% PBST for 2 hours at room temperature. Membranes were washed again 3x with 0.1% PBST before being incubated with a 1:1 solution of Pierce ECL Western Blotting Substrate (Thermo Scientific). Chemiluminescence was detected on a ChemiDoc™ MP Imaging System darkroom (BioRad). Densitometric readings of band intensities for blots were obtained using ImageJ™. Primary antibodies (all used at 1:1000 dilution and purchased from Cell Signalling Technology, unless noted otherwise) were: anti-NRP2 (clone D39A5), anti-HSC70 (clone B-6; Santa Cruz Biotechnology), anti-p-VASP^Ser157^ (clone 3111), anti-VASP (clone 3132), anti-p-FAK^Tyr407^ (clone 44-650G; Invitrogen), anti-tensin-1 (clone NBP1-84129; Novus Biologicals), anti-GAPDH (60004-1-1g; Proteintech), EDA-FN (F6140; Sigma).

### Signalling Assays

siRNA-transfected ECs were seeded into FN-coated 6 cm cultures dishes at a density of 5×10^5^ cells/well and incubated for 48 hours. ECs were then PBS washed and starved for 3 hours in serum free medium (OptiMEM^^®^^; Invitrogen). VEGF was then added at a final concentration of 30 ng/ml at indicated timepoints. ECs were then subjected to lysing, protein quantification and protein expression analysis by Western blot.

### Co-Immunoprecipitation Assays

siRNA-transfected ECs were seeded into FN-coated 10 cm dishes at a density of 2×10^6^ cells/dish, and incubated for 48 hours. ECs were then lysed on ice in lysis buffer as previously described by Valdembri et al (Valdembri *et al.*, 2009) in the presence of 100X Halt protease inhibitor cocktail (Thermo Scientific) and protein quantified using the DC BioRad assay. 200 μg protein from each sample was immunoprecipitated by incubation with protein-G Dynabeads^^®^^ (Invitrogen) coupled to a rabbit anti-α5-integrin antibody (clone 4705S, Cell Signalling Technology) on a rotator overnight at 4°C. Immunoprecipitated complexes were then washed 3x with lysis buffer + 100X Halt™ protease inhibitor, and once in PBS, before being added to and boiled in NuPAGE sample reducing agent and sample buffer (Life Technologies) for Western blot analysis.

### Deoxycholate (DOC) Buffer-extraction

siRNA-treated ECs were allowed to adhere for 16 hours at 37°C and 5% CO_2._ ECs were then lysed in DOC lysis buffer (20mM Tris, pH 8.5, 1% sodium deoxycholate, 2mM iodoacetamide, 2mM EDTA) in the presence of 100X Halt protease inhibitor cocktail, cleared by centrifugation, and the insoluble fraction isolated. Insoluble fractions were separated by SDS-PAGE and subjected to Western blot analysis.

### *In vivo* CMT19T Tumour Growth Assays

Inducible, endothelial specific deletion of NRP2 was achieved by crossing mice expressing the PDGFb.iCreER promoter of Cre-recombinase to those floxed for NRP2. NRP2^*flfl*^; PDGFb.iCreER and NRP2^*flfl*^ littermate control mice received intraperitoneal (IP) injections of tamoxifen (75 mg/kg bodyweight, 2mg/ml stock) thrice weekly (Monday, Wednesday, Friday) for the duration of the experiment from D-4 to D17 to induce Cre-recombinase activity. CMT19T lung carcinoma cells (CR-UK Cell Production) (1×10^6^) were implanted subcutaneously (SC) into the flank of mice at D0 and allowed to grow until D18. On D18, mice were killed, tumour sizes measured, and tumour samples fixed in 4% PFA for blood vessel density analysis. Tumour volumes were calculated according to the formula: length x width^2^ x 0.52.

### Immunofluorescence Analysis of Tumour Sections

Tumour sections were fixed in 4% PFA for 10 minutes at RT before being washed twice in PBS 0.3% triton-X100, twice in PBLEC (1x PBS, 1% Tween 20, 0.1 mM CaCl_2,_ 0.1 mM MgCl_2,_ 0.1 mM MnCl_2_) and incubated in Dako protein block serum free (Agilent). Sections were then incubated overnight at 4°C in primary antibodies against NRP2 (clone Sc-13117; Santa-Cruz Biotechnology) and endomucin (clone Sc-65495; Santa-Cruz Biotechnology). Following primary antibody incubation, sections were washed again twice in PBS 0.3% triton-X100 and PBLEC before being incubated in the appropriate Alexa fluor secondary antibody for 2 hours at RT. Sections were then blocked in Sudan black for 5 minutes before being mounted with flouromount + DAPI and imaged at 10 X magnification using a Zeiss AxioImager M2 microscope (AxioCam MRm camera). Blood-vessel density was assessed by counting the number of endomucin-positive vessels per mm^2^ from 3 representative ROIs/ section, averaged over 3 sections/tumour.

### *In vivo* Retina Assays

Inducible, endothelial specific deletion of NRP2 was achieved by crossing mice expressing the PDGFb.iCreER promoter of Cre-recombinase to those floxed for NRP2. Tamoxifen-induced activation of Cre-recombinase and thus deletion of NRP2 was employed via 2 regimes: NRP2^*flfl*^; PDGFb.iCreER and NRP2^*flfl*^ littermate control mice either received subcutaneous (SC) injections of tamoxifen (50 μl, 2mg/ml stock) on postnatal (P) days 2 and 3, followed by intraperitoneal (IP) injections of the same dose on P4 and 5 before retinas being harvested at P6, or mice received IP injections of tamoxifen (50 μl, 2mg/ml stock) on postnatal (P) days 7 to 10 before retinas being harvested at P12. After dissection, retinas were fixed in ice-cold methanol for 30 minutes before being prepared for blocking and immuno-staining. First, retinas were permeabilised in PBS 0.25% triton-X100 for 30 minutes at RT, before being washed twice in PBLEC and then blocked in Dako protein block serum free for 1 hour. Retinas were then incubated in primary antibodies against: FITC-BS-1 lectin (clone L2895; Sigma), NRP2 (clone Sc-13117; Santa-Cruz Biotechnology), αSMA (clone Ab21027; Abcam), ERG 1 / 2 (clone Ab92513; Abcam), p-FAK^Tyr407^ (clone 44-650G Invitrogen), Collagen IV (clone Ab19808; Abcam). Following primary antibody incubation, retinas were incubated in the appropriate Alexa fluor secondary antibody for 1 hour at RT, before being washed twice in PBS 0.1% triton-X100 and mounted. Images were captured using a Zeiss LSM880 Airyscan Confocal microscope. Vascular extension analysis was performed using ImageJ™, vessel density analysis was performed using AngioTool™.

### Statistical Analysis

The graphic illustrations and analyses to determine statistical significance were generated using GraphPad Prism 6 software and Student’s t-tests, respectively. Bar charts show mean values and the standard error of the mean (+SEM). Asterisks indicate the statistical significance of P values: P > 0.05 = NS (not significant), * P < 0.05, ** P < 0.01, *** P < 0.001 and **** P < 0.0001.

## Author Contributions

Conceptualisation: CJB, JAGET, SDR; Formal analyses: CJB, JAGET, SDR; Investigation: CJB, JAGET, SDR; Resources: SDR; Review and editing: CJB, JAGET, SDR; Visualisation: CJB, JAGET, SDR; Supervision: SDR; Funding acquisition: SDR.

## Acknowledgments

This work was supported by funding from: the UKRI Biotechnology and Biological Sciences Research Council Norwich Research Park Biosciences Doctoral Training Partnership and BHF (grant number PG/15/25/31369); Additionally, we thank Norfolk Fundraisers, Mrs Margaret Doggett, and the Colin Wright Fund for their kind support and fundraising over the years. Robinson is also partially funded by the BBSRC Institute Strategic Programme Gut Microbes and Health BB/R012490/1 and its constituent project (BBS/E/F/000PR10355).

## Data availability statement

The raw data supporting the conclusions of this manuscript will be made available by the authors, without undue reservation, to any qualified researcher.

## References

Ajeian, J. N. et al. (2016) ‘Proteomic analysis of integrin-associated complexes from mesenchymal stem cells’, Proteomics - Clinical Applications, 10(1), pp. 51–57. doi: 10.1002/prca.201500033.

Alarcon-Martinez, L. et al. (2018) ‘Capillary pericytes express α-smooth muscle actin, which requires prevention of filamentous-actin depolymerization for detection’, eLife, 7, pp. 1–17. doi: 10.7554/eLife.34861.

Alghamdi, A. A. A. et al. (2020) ‘NRP2 as an Emerging Angiogenic Player; Promoting Endothelial Cell Adhesion and Migration by Regulating Recycling of α5 Integrin’, Frontiers in Cell and Developmental Biology, 8(May), pp. 1–16. doi: 10.3389/fcell.2020.00395.

Bachelder, R. E. et al. (2001) ‘Vascular endothelial growth factor is an autocrine survival factor for neuropilin-expressing breast carcinoma cells’, Cancer Research, 61(15), pp. 5736–5740.

Benz, P. M. et al. (2009) ‘Differential VASP phosphorylation controls remodeling of the actin cytoskeleton’, Journal of Cell Science, 122(21), pp. 3954–3965. doi: 10.1242/jcs.044537.

Bernusso, V. A. et al. (2015) ‘Imatinib restores VASP activity and its interaction with Zyxin in BCR-ABL leukemic cells’, Biochimica et Biophysica Acta - Molecular Cell Research. Elsevier B.V., 1853(2), pp. 388–395. doi: 10.1016/j.bbamcr.2014.11.008.

Bielenberg, D. R. et al. (2004) ‘Semaphorin 3F, a chemorepulsant for endothelial cells, induces a …’, J Clin Invest, 114(9), pp. 1260–1271. doi: 10.1172/JCI200421378.1260.

Bielenberg, D. R. et al. (2006) ‘Neuropilins in neoplasms: Expression, regulation, and function’, Experimental Cell Research, 312(5), pp. 584–593. doi: 10.1016/j.yexcr.2005.11.024.

Bielenberg, D. R. et al. (2012) ‘Increased smooth muscle contractility in mice deficient for neuropilin 2’, American Journal of Pathology. Elsevier Inc., 181(2), pp. 548–559. doi: 10.1016/j.ajpath.2012.04.013.

Bökel, C. and Brown, N. H. (2002) ‘Integrins in development: Moving on, responding to, and sticking to the extracellular matrix’, Developmental Cell, 3(3), pp. 311–321. doi: 10.1016/S1534-5807(02)00265-4.

Borkowetz, A. et al. (2020) ‘Neuropilin-2 is an independent prognostic factor for shorter cancer-specific survival in patients with acinar adenocarcinoma of the prostate’, International Journal of Cancer, 146(9), pp. 2619–2627. doi: 10.1002/ijc.32679.

Brash, J. T. et al. (2020) ‘Tamoxifen-Activated CreERT Impairs Retinal Angiogenesis Independently of Gene Deletion’, pp. 849–850. doi: 10.1006/bbrc.1997.7124.

Buskermolen, A. B. C., Kurniawan, N. A. and Bouten, C. V. C. (2018) ‘An automated quantitative analysis of cell, nucleus and focal adhesion morphology’, PLoS ONE, 13(3), pp. 1–16. doi: 10.1371/journal.pone.0195201.

Calderwood, D. A. (2004) ‘Integrin activation’, Journal of Cell Science, 117(5), pp. 657–666. doi: 10.1242/jcs.01014.

Cao, Y. et al. (2013) ‘Neuropilin-2 promotes extravasation and metastasis by interacting with endothelial α5 integrin’, Cancer Research, 73(14), pp. 4579–4590. doi: 10.1158/0008-5472.CAN-13-0529.

Chang, F. et al. (2007) ‘FAK Potentiates Rac1 Activation and Localization to Matrix Adhesion Sites: A Role for Beta-PIX’, Molecular Biology of the Cell, 18, pp. 253–264.

Clark, R. A. F. et al. (1982) ‘Blood vessel fibronectin increases in conjunction with endothelial cell proliferation and capillary ingrowth during wound healing’, Journal of Investigative Dermatology, 79(5), pp. 269–276. doi: 10.1111/1523-1747.ep12500076.

Deshpande, P. P., Biswas, S. and Torchilin, V. P. (2013) ‘Current trends in the use of liposomes for tumor targeting’, Nanomedicine, 8(9), pp. 1509–1528. doi: 10.2217/nnm.13.118.

Ellison, T. S. et al. (2015) ‘Suppression of β3-integrin in mice triggers a neuropilin-1-dependent change in focal adhesion remodelling that can be targeted to block pathological angiogenesis’, DMM Disease Models and Mechanisms, 8(9), pp. 1105–1119. doi: 10.1242/dmm.019927.

Fakhari, M. et al. (2002) ‘Selective upregulation of vascular endothelial growth factor receptors neuropilin-1 and −2 in human neuroblastoma’, Cancer, 94(1), pp. 258–263. doi: 10.1002/cncr.10177.

Fantin, A. et al. (2011) ‘The cytoplasmic domain of neuropilin 1 is dispensable for angiogenesis, but promotes the spatial separation of retinal arteries and veins’, Development, 138(19), pp. 4185–4191. doi: 10.1242/dev.070037.

Fantin, A. et al. (2015) ‘NRP1 Regulates CDC42 Activation to Promote Filopodia Formation in Endothelial Tip Cells’, Cell Reports. The Authors, 11(10), pp. 1577–1590. doi: 10.1016/j.celrep.2015.05.018.

Favier, B. et al. (2006) ‘Neuropilin-2 interacts with VEGFR-2 and VEGFR-3 and promotes human endothelial cell survival and migration’, Blood, 108(4), pp. 1243–1250. doi: 10.1182/blood-2005-11-4447.

Fukahi, K. et al. (2004) ‘Aberrant Expression of Neuropilin-1 and −2 in Human Pancreatic Cancer Cells’, Clinical Cancer Research, 10(2), pp. 581–590. doi: 10.1158/1078-0432.CCR-0930-03.

Gao, C. et al. (2015) ‘FAK/PYK2 promotes the Wnt/β-catenin pathway and intestinal tumorigenesis by phosphorylating GSK3β’, eLife, 4(AUGUST2015), pp. 1–17. doi: 10.7554/eLife.10072.

Ghosh Dastidar, D., Ghosh, D. and Chakrabarti, G. (2020) ‘Tumour vasculature targeted anti-cancer therapy’, Vessel Plus, 2020. doi: 10.20517/2574-1209.2019.36.

Goel, H. L. et al. (2011) ‘Neuropilin-2 promotes branching morphogenesis in the mouse mammary gland’, Development, 138(14), pp. 2969–2976. doi: 10.1242/dev.051318.

Goel, H. L. et al. (2012) ‘Neuropilin-2 regulates α6β1 integrin in the formation of focal adhesions and signaling’, Journal of Cell Science, 125(2), pp. 497–506. doi: 10.1242/jcs.094433.

Goley, E. and Welch, M. (2006) ‘The ARP2/3 complex: an actin nucleator comes of age’, Nature Reviews Molecular Cell Biology, 7, pp. 713–726.

Golubovskaya, V. M. et al. (2009) ‘The direct effect of Focal Adhesion Kinase (FAK), dominant-negative FAK, FAK-CD and FAK siRNA on gene expression and human MCF-7 breast cancer cell tumorigenesis’, BMC Cancer, 9. doi: 10.1186/1471-2407-9-280.

Hanahan, D. and Weinberg, R. A. (2011) ‘Hallmarks of cancer: The next generation’, Cell. Elsevier Inc., 144(5), pp. 646–674. doi: 10.1016/j.cell.2011.02.013.

Herzog, B. et al. (2011) ‘VEGF binding to NRP1 is essential for VEGF stimulation of endothelial cell migration, complex formation between NRP1 and VEGFR2, and signaling via FAK Tyr407 phosphorylation’, Molecular Biology of the Cell, 22(15), pp. 2766–2776. doi: 10.1091/mbc.E09-12-1061.

Holt, M. R., Critchley, D. R. and Brindle, N. P. J. (1998) ‘The focal adhesion phosphoprotein, VASP’, International Journal of Biochemistry and Cell Biology, 30(3), pp. 307–311. doi: 10.1016/S1357-2725(97)00101-5.

Hu, P. and Luo, B. H. (2013) ‘Integrin bi-directional signaling across the plasma membrane’, Journal of Cellular Physiology, 228(2), pp. 306–312. doi: 10.1002/jcp.24154.

Hynes, R. (2002) ‘Integrins: Bidirectional Allosteric Signaling Machines’, Cell, 110, pp. 673–687. Available at: https://www.ncbi.nlm.nih.gov/pubmed/12297042.

Ingber, D. E. (1990) ‘Fibronectin controls capillary endothelial cell growth by modulating cell shape’, Proceedings of the National Academy of Sciences of the United States of America, 87(9), pp. 3579–3583. doi: 10.1073/pnas.87.9.3579.

Jalal, S. et al. (2019) ‘Actin cytoskeleton self-organization in single epithelial cells and fibroblasts under isotropic confinement’, Journal of Cell Science, 132(5), pp. 1–14. doi: 10.1242/jcs.220780.

Jianliang Zhang and Steven N. Hochwald (2014) ‘The role of FAK in tumor metabolism and therapy’, Pharmacol Ther, 142(2), pp. 154–163. doi: 10.1016/j.pharmthera.2013.12.003.

Kaksonen, M., Toret, C. P. and Drubin, D. G. (2006) ‘Harnessing actin dynamics for clathrin-mediated endocytosis’, Nature Reviews Molecular Cell Biology, 7(6), pp. 404–414. doi: 10.1038/nrm1940.

Kawakami, T. et al. (2002) ‘Neuropilin 1 and neuropilin 2 co-expression is significantly correlated with increased vascularity and poor prognosis in nonsmall cell lung carcinoma’, Cancer, 95(10), pp. 2196–2201. doi: 10.1002/cncr.10936.

Kragtorp, K. A. and Miller, J. R. (2006) ‘Regulation of somitogenesis by Ena/VASP proteins and FAK during Xenopus development’, Development, 133(4), pp. 685–695. doi: 10.1242/dev.02230.

Krilleke, D. et al. (2007) ‘Molecular mapping and functional characterization of the VEGF164 heparin-binding domain’, Journal of Biological Chemistry, 282(38), pp. 28045–28056. doi: 10.1074/jbc.M700319200.

Lambert, J. et al. (2020) ‘ADAMTS-1 and syndecan-4 intersect in the regulation of cell migration and angiogenesis’, Journal of Cell Science, 133(7), pp. 1–15. doi: 10.1242/jcs.235762.

Lantuéjoul, S. et al. (2003) ‘Expression of VEGF, semaphorin SEMA3F, and their common receptors neuropilins NP1 and NP2 in preinvasive bronchial lesions, lung tumours, and cell lines’, Journal of Pathology, 200(3), pp. 336–347. doi: 10.1002/path.1367.

Lenzo, F. L. and Cance, W. G. (2017) ‘From tumorigenesis to microenvironment and immunoregulation: The many faces of focal adhesion kinase and challenges associated with targeting this elusive protein’, Translational Cancer Research, 6(17), pp. S957–S960. doi: 10.21037/tcr.2017.06.05.

Luo, X. et al. (2020) ‘Vascular NRP2 triggers PNET angiogenesis by activating the SSH1-cofilin axis’, Cell and Bioscience. BioMed Central, 10(1), pp. 1–16. doi: 10.1186/s13578-020-00472-6.

Mana, G. et al. (2016) ‘PPFIA1 drives active α5β1 integrin recycling and controls fibronectin fibrillogenesis and vascular morphogenesis’, Nature Communications, 7(November). doi: 10.1038/ncomms13546.

Michael, K. E. et al. (2009) ‘Focal adhesion kinase modulates cell adhesion strengthening via integrin activation’, Molecular Biology of the Cell, 20(9), pp. 2508–2519. doi: 10.1091/mbc.E08-01-0076.

Milde, F. et al. (2013) ‘The mouse retina in 3D: Quantification of vascular growth and remodeling’, Integrative Biology (United Kingdom), 5(12), pp. 1426–1438. doi: 10.1039/c3ib40085a.

Nader, G. P. F., Ezratty, E. J. and Gundersen, G. G. (2016) ‘FAK, talin and PIPKI 3 regulate endocytosed integrin activation to polarize focal adhesion assembly’, Nature Cell Biology, 18(5), pp. 491–503. doi: 10.1038/ncb3333.

Nagano, M. et al. (2012) ‘Turnover of focal adhesions and cancer cell migration’, International Journal of Cell Biology, (June 2014). doi: 10.1155/2012/310616.

Pan, L. et al. (2016) ‘Research advances on structure and biological functions of integrins’, SpringerPlus. Springer International Publishing, 5(1). doi: 10.1186/s40064-016-2502-0.

De Pascalis, C. and Etienne-Manneville, S. (2017) ‘Single and collective cell migration: The mechanics of adhesions’, Molecular Biology of the Cell, 28(14), pp. 1833–1846. doi: 10.1091/mbc.E17-03-0134.

Plein, A., Fantin, A. and Ruhrberg, C. (2014) ‘Neuropilin regulation of angiogenesis, arteriogenesis, and vascular permeability’, Microcirculation, 21(4), pp. 315–323. doi: 10.1111/micc.12124.

Raimondi, C. et al. (2014) ‘Imatinib inhibits VEGF-independent angiogenesis by targeting neuropilin 1-dependent ABL1 activation in endothelial cells’, Journal of Experimental Medicine, 211(6), pp. 1167–1183. doi: 10.1084/jem.20132330.

Reynolds, L. and Hodivala-Dilke, K. (2006) ‘Primary mouse endothelial cell culture for assays of angiogenesis.’, Methods Mol Med, 120, pp. 503–509.

Robinson, S. D. et al. (2009) ‘Αvβ3 Integrin Limits the Contribution of Neuropilin-1 To Vascular Endothelial Growth Factor-Induced Angiogenesis’, Journal of Biological Chemistry, 284(49), pp. 33966–33981. doi: 10.1074/jbc.M109.030700.

Robinson, S. D. and Hodivala-Dilke, K. M. (2011) ‘The role of β3-integrins in tumor angiogenesis: Context is everything’, Current Opinion in Cell Biology. Elsevier Ltd, 23(5), pp. 630–637. doi: 10.1016/j.ceb.2011.03.014.

Rossier, O. et al. (2012) ‘Integrins β 1 and β 3 exhibit distinct dynamic nanoscale organizations inside focal adhesions’, Nature Cell Biology, 14(10), pp. 1057–1067. doi: 10.1038/ncb2588.

Sadok, A. and Marshall, C. J. (2014) ‘Rho gtpases masters of cell migration’, Small GTPases, 5(4). doi: 10.4161/sgtp.29710.

Schaller, M. D. (2010) ‘Cellular functions of FAK kinases: Insight into molecular mechanisms and novel functions’, Journal of Cell Science, 123(7), pp. 1007–1013. doi: 10.1242/jcs.045112.

Schimmel, L. et al. (2020) ‘C-Src controls stability of sprouting blood vessels in the developing retina independently of cell-cell adhesion through focal adhesion assembly’, Development (Cambridge), 147(7). doi: 10.1242/dev.185405.

Srichai, M. and Zent, R. (2010) Integrin Structure and Function: Cell-Extracellular Matrix Interactions in Cancer. Springer, Boston, MA.

Sundararaman, A. et al. (2020) ‘RhoJ Regulates α5β1 Integrin Trafficking to Control Fibronectin Remodeling during Angiogenesis’, Current Biology. Elsevier Ltd., 30(11), pp. 2146–2155.e5. doi: 10.1016/j.cub.2020.03.042.

Uruno, T. et al. (2001) ‘Activation of Arp2/3 complex-mediated actin polymerization by cortactin’, Nature Cell Biology, 3(3), pp. 259–266. doi: 10.1038/35060051.

Valdembri, D. et al. (2009) ‘Neuropilin-1/GIPC1 signaling regulates α5β1 integrin traffic and function in endothelial cells’, PLoS Biology, 7(1). doi: 10.1371/journal.pbio.1000025.

Vales, A. et al. (2007) ‘Myeloid leukemias express a broad spectrum of VEGF receptors including neuropilin-1 (NRP-1) and NRP-2’, Leukemia and Lymphoma, 48(10), pp. 1997–2007. doi: 10.1080/10428190701534424.

Woo, M. K. and Fowler, V. M. (1994) ‘Identification and characterization of tropomodulin and tropomyosin in the adult rat lens’, Journal of Cell Science, 107(5), pp. 1359–1367.

Woodside, D., Liu, S. and Ginsberg, M. (2001) ‘Integrin activation’, Thromb Haemost, 86, pp. 316–323.

Yamakita, Y. et al. (1996) ‘Phosphorylation of human fascin inhibits its actin binding and bundling activities’, Journal of Biological Chemistry, 271(21), pp. 12632–12638. doi: 10.1074/jbc.271.21.12632.

Zamir, E. et al. (2000) ‘Dynamics and segregation of cell-matrix adhesions in cultured fibroblasts’, Nature Cell Biology, 2(4), pp. 191–196. doi: 10.1038/35008607.

